# Specialized germline P-bodies are required to specify germ cell fate in *C. elegans* embryos

**DOI:** 10.1101/2022.08.15.504042

**Authors:** Madeline Cassani, Geraldine Seydoux

## Abstract

In animals with germ plasm, specification of the germline involves “germ granules”, cytoplasmic condensates that enrich maternal transcripts in the germline founder cells. In *C. elegans* embryos, P granules enrich maternal transcripts, but surprisingly P granules are not essential for germ cell fate specification. Here we describe a second condensate in the *C. elegans* germ plasm. Like canonical P-bodies found in somatic cells, “germline P-bodies” contain regulators of mRNA decapping and deadenylation and, in addition, the intrinsically-disordered proteins MEG-1 and MEG-2 and the TIS11-family RNA-binding protein POS-1. Embryos lacking *meg-1* and *meg-2* do not stabilize P-body components, miss-regulate POS-1 targets, miss-specify the germline founder cell, and do not develop a germline. Our findings suggest that specification of the germ line involves at least two distinct condensates that independently enrich and regulate maternal mRNAs in the germline founder cells.

## Introduction

The germ plasm is a specialized cytoplasm found in the eggs of certain insects, nematodes and vertebrates that serves as a vehicle to segregate maternal proteins and RNAs to the nascent embryonic germline (Kulkarni and Extavour, 2017). Germ plasm assembly is a derived trait that arose independently several times in evolution as an alternative to the ancestral mode of germ cell fate specification by cell-to-cell signaling (Kemph and Lynch, 2022). A convergent characteristic of germ plasm in both vertebrate and invertebrate species is the presence of “germ granules”, micron-size ribonucleoprotein assemblies that contain RNAs coding for factors that promote germ cell development (Kulkarni and Extavour, 2017). Germ granules segregate with the germ plasm to the germline founder cells and are thought to contribute to their specification as primordial germ cells (“PGCs”)· Germ granules were initially described using electron microscopy as mostly amorphous, electron-dense, micron-sized structures not surrounded by membranes (Arkov and Ramos, 2010). Fluorescence microscopy studies and proteomics in *Drosophila*, zebrafish, *Xenopus, C. elegans* and mice have revealed the presence of different types of condensates in germ cells, some with complex sub-structure (Gallo et al., 2008; Wang et al., 2014; Vo et al., 2019; Eichler et al., 2020; Wan et al., 2018; Uebel et al., 2020; Uebel et al., 2021; Roovers et al., 2018; Neil et al., 2021; Yang et al., 2022; Aravin et al., 2009). These studies have hinted that germ cells contain multiple condensates that compartmentalize different RNA-centered activities that collectively specify germ cell fate. For example, polar granules and founder granules are distinct granules in the germ plasm of *Drosophila melanogaster* that harbor mRNAs that need to be translated (polar granules) or degraded (founder granules) for proper germline development (Eichler et al., 2020). Here we demonstrate that the *C. elegans* germ plasm also contains two condensate types that make distinct contributions towards germ cell fate.

The first condensates to be described in the *C. elegans* germ plasm were named P granules for their segregation with P (posterior) blastomeres through a series of 4 asymmetric divisions that eventually give rise to the germline founder cell P_4_ (Strome and Wood, 1982; Fig. 1E). P granules are scaffolded by the nematode-specific, RGG-domain proteins PGL-1 and PGL-3, which form dense liquid-like condensates *in vitro* and *in vivo* (Brangwynne et al., 2009; Hanazawa et al., 2011; Updike et al., 2011; Saha et al., 2016; Putnam et al., 2019). In zygotes, the PGL condensates become covered on their surface by nanoscale solid clusters assembled by a pair of paralogous and redundant intrinsically-disordered proteins MEG-3 and MEG-4. MEG-3/4 form an essential protective layer that controls the dynamics and asymmetric segregation of PGL condensates into the P blastomeres in part by reducing the surface tension of PGL condensates (Folkmann and Putnam, et al., 2021). MEG-3/4 also recruit maternal mRNAs to P granules. MEG-3 binds RNA *in vitro* and co-precipitates with ~500 maternal mRNAs in embryonic lysates, including the Nanos homologue *nos-2* and the predicted E3 ubiquitin ligase *Y51F10.2* that are required redundantly for fertility (Lee et al., 2020). Incorporation into P granules enriches RNAs in the P_4_ blastomere as much as 5-fold over what would have been achieved by equal segregation to all embryonic cells (Schmidt et al., 2021). *nos-2* and *Y51F10.2* are translationally repressed in the P_0_ to P_3_ blastomeres and become translationally activated in P_4_, the germline founder cell (Subramaniam and Seydoux, 1999; Lee et al., 2020). Despite their role in enriching mRNAs required for germ cell development, P granules are not essential for germ cell fate. In *meg-3 meg-4* mutants, the germline founder cell P_4_ inherits no PGL condensates and reduced levels of *nos-2* and *Y51F10.2* transcripts (Lee et al., 2020; Schmidt et al., 2021). These transcripts, however, are still translationally activated in P_4_, and *meg-3 meg-4* animals are mostly (~70%) fertile (Lee et al., 2020). These observations indicate that the *C. elegans* germ plasm maintains proper regulation of maternal mRNAs in the absence of P granules.

**Fig. 1:**
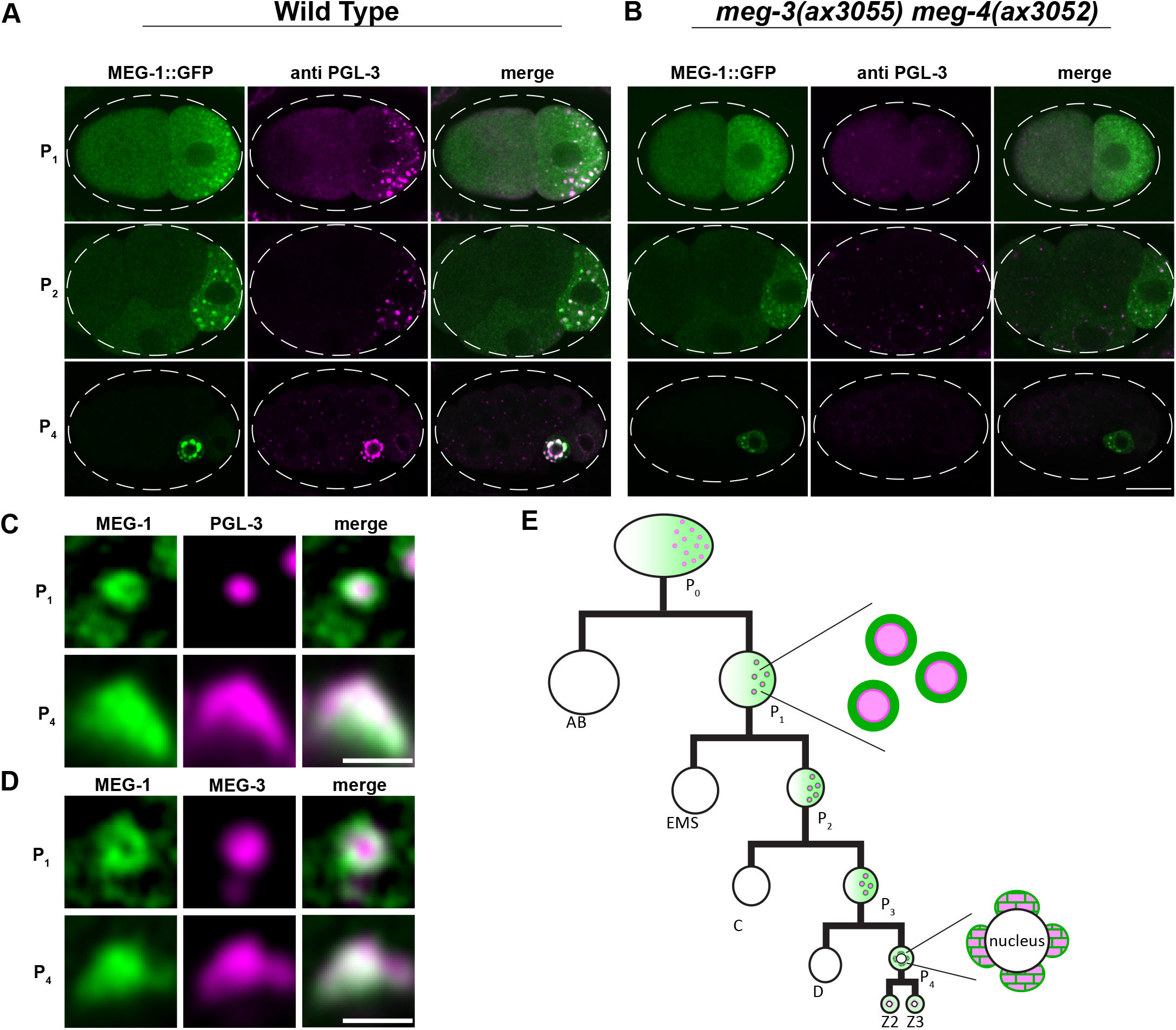
MEG-1 puncta are distinct from P granules. Representative Airyscan photomicrographs of wild-type (A) and *meg-3 meg-4* mutant (B) embryos expressing endogenous MEG-1::GFP and co-stained for GFP and PGL-3. MEG-1, but not PGL-3, enriches in P blastomeres in *meg-3 meg-4* embryos. Scale bar is 10 μm. (C and D) Higher resolution images of MEG-1::GFP and PGL-3 (C) and MEG-1::GFP and MEG-3::OLLAS (D) in P_1_ and P_4_. In P_1_, MEG-1 enriches at the periphery of PGL-3 and MEG-3. In P_4_, P granules become perinuclear and MEG-1 and PGL-3/MEG-3 overlap. See Fig. S1A for quantification. Scale bars are 1 μm. (E) Abbreviated cartoon lineage summarizing the distribution of MEG-1 (green) and P granules (pink) in the germline (P) blastomeres. In the zygote P_0_, MEG-1 is present in a cytoplasmic gradient as well as small granules that are difficult to visualize at this stage. MEG-1 enriches at the periphery of P granules in the P_1-3_ blastomeres, and merges with P granules in P_4_. In the primordial germ cells Z2 and Z3, MEG-1 becomes cytoplasmic and is degraded, while P granules remain.

The *C. elegans* germ plasm contains a second condensate type that contains proteins characteristic of P-bodies, ubiquitous RNP granules implicated in mRNA storage and decay (Gallo et al., 2008; Ivanov et al., 2019). P-body-like condensates associate with P granules in germ plasm in tight assemblies containing a central P granule surrounded by several P-body-like condensates (Gallo et al., 2008). Dozens of proteins have been reported to enrich in granules in the *C. elegans* germ plasm (Updike and Strome, 2010; Phillips and Updike, 2022), and, in most cases, it is not known whether these localize to P granules proper (as defined by PGL-3 and MEG-3) or to the closely apposed P-body-like condensates described in Gallo et al. 2008, or to both. In particular, MEG-1 and MEG-2 are two intrinsically-disordered proteins, distantly related to MEG-3 and MEG-4, and originally described as P granule proteins (Leacock and Reinke, 2008). In this study, we demonstrate that MEG-1 and MEG-2 associate with canonical P-body proteins and stabilize P-body-like condensates in P_4_. Our findings indicate that, unlike P granules, “germline P-bodies” are essential for maternal mRNA regulation and specification of P_4_ as the germline founder cell.

## Results

### MEG-1 enriches in puncta distinct from P granules

To characterize the localization of MEG-1, we used a MEG-1::GFP fusion where GFP is inserted at the C-terminus of the MEG-1 ORF in the *meg-1* locus. Consistent with a previous report that used a polyclonal antibody raised against MEG-1 (Leacock and Reinke, 2008), MEG-1::GFP segregated with germ plasm in early embryos, distributing between a cytoplasmic pool and bright puncta in P blastomeres that overlapped with P granules (Fig. 1A). High resolution images revealed that the MEG-1 puncta localize to the periphery of P granules (visualized with PGL-3 or MEG-3) in P_1_ blastomeres (Fig. 1C,D, Fig. S1A). By the P_4_ stage, when P granules are fully perinuclear, the MEG-1::GFP signal was distributed throughout P granules (Fig. 1C,D, Fig. S1A). In Z2 and Z3, MEG-1::GFP dispersed back into the cytoplasm (Fig. S1B) and turned over by mid-embryogenesis (Leacock and Reinke, 2008).

Leacock and Reinke, 2008 reported that MEG-1 enrichment in P blastomeres is independent of P granule components and vice versa. Consistent with these results, we found that MEG-1 still enriched preferentially in P blastomeres in *meg-3(ax3055) meg-4(ax3052)* mutants (Fig. 1B). MEG-1 puncta, however, remained cytoplasmic and did not associate with the nuclear envelope in P_4_ of *meg-3(ax3055) meg-4(ax3052)* mutants (Fig. 1A-B). Leacock and Reinke, 2008 used a partial deletion of the *meg-1* locus and RNAi of the *meg-1* paralog *meg-2* to generate embryos depleted of both *meg-1* and *meg-2*. To complement these analyses, we created a deletion that removed the entire *meg-1 meg-2* operon. *meg-1 meg-2(ax4532)* hermaphrodites were 100% maternal effect sterile as reported for *meg-1(vr10) meg-2(RNAi)* (Table S1). We found that MEG-3 and PGL-3 still assembled into puncta that segregated with P blastomeres in *meg-1 meg-2(ax4532)* embryos, confirming that P granule assembly does not require *meg-1* and *meg-2* (Fig. S1C). We noticed, however, that P granule enrichment in P blastomeres was not as robust in *meg-1 meg-2* embryos (Fig. S1D) as previously reported (Leacock and Reinke, 2008; Wang et al., 2014), suggesting a minor contribution of MEG-1/2 to P granule segregation.

We conclude that MEG-1 localizes to assemblies that are distinct from P granules. MEG-1 puncta and P granules interact but assemble independently in the cytoplasm of P blastomeres.

### MEG-1 immunoprecipitates with P-body components and several RNA-binding proteins, including POS-1

As we show here for MEG-1, we previously reported that P-body markers enrich at the periphery of P granules in early P blastomeres (Gallo et al., 2008). Furthermore, Wu et al., 2017 identified MEG-1 and MEG-2 among immunoprecipitates of the P-body scaffold C-NOT1^NTL-1^ and identified seven CCR4-NOT subunits in MEG-2 immunoprecipitates. To complement these studies, we performed mass spectrometry on MEG-1::GFP immunoprecipitated from early embryo lysates using anti-GFP antibodies. As controls, we used lysates from wild-type worms expressing untagged MEG-1. We identified 54 proteins that were enriched at least twofold over untagged controls in two biological replicates (Fig. 2A, Table S2).

**Fig. 2:**
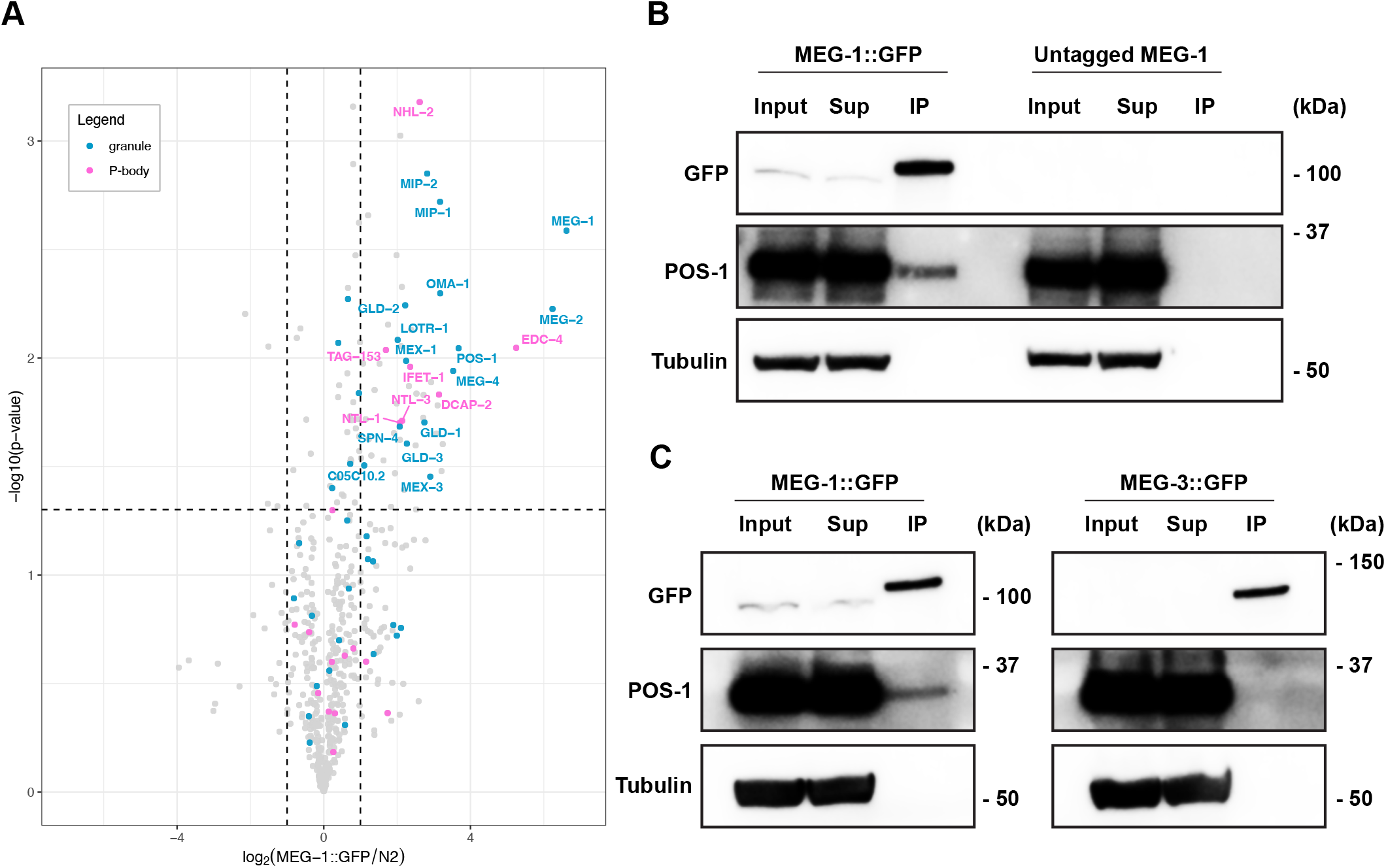
MEG-1 immunoprecipitates with P-body and RNA-binding proteins, including POS-1. (A) Volcano plot showing on the X-axis the log2 fold enrichment of proteins (dots) in MEG-1::GFP immunoprecipitates over “N2” (wild-type lysates containing untagged MEG-1) as a function of the log10 *p*-value calculated from two independent immunoprecipitation experiments (Y-axis). Of the 54 proteins enriched in MEG-1::GFP immunoprecipitates (top right quadrant), 13% correspond to P-body proteins (labeled in pink) and 28% correspond to proteins previously reported to localize to granules in P blastomeres (blue). (B) Representative western blots from two independent experiments confirm that GFP immunoprecipitates pull down MEG-1::GFP and POS-1, but not tubulin. (C) Western blots from MEG-1::GFP and MEG-3::GFP immunoprecipitates. Unlike MEG-1::GFP, MEG-3::GFP does not pull down POS-1. Full western blot images are shown in Fig. S2.

Among the proteins in MEG-1::GFP immunoprecipitates, we observed an enrichment for canonical P-body proteins (7 out of 36 canonical P-body proteins in the *C. elegans* genome/WormBase, p<0.0001, Fisher’s exact test), including the decapping factors DCP2^DCAP-2^ and EDC4^EDC-4^, the TRIM-NHL family member and miRISC cofactor TRIM45^NHL-2^, the CCR4-NOT complex subunits CNOT1^NTL-1^, CNOT2^TAG-153^, CNOT3^NTL-3^ and the translational repressor and DDX6-binding partner eIF4-ET^IFET-1^ (Table S2). In addition to P-body proteins, we also observed eight RNA-binding proteins including the translational repressor GLD-1, the poly-A polymerase GLD-2/GLD-3, the zinc finger proteins MEX-1, OMA-1 and POS-1, the KH domain protein MEX-3, and the RRM domain protein SPN-4. All of these have been reported to regulate maternal mRNAs and to enrich in germ plasm and “P granules” (because P-bodies and P granules are closely linked in wild-type embryos, most studies have not distinguished between the two). Among these, POS-1 scored as one of the most highly enriched proteins in MEG-1::GFP precipitates after MEG-1 and MEG-2 (Fig. 2A, Table S2).

POS-1 regulates the poly-adenylation of thousands of maternal mRNAs containing AU-rich elements (AREs) in their 3’UTR (Farley et al., 2008; Elewa et al., 2015). ARE-binding proteins have been reported to recruit P-body components, including decapping enzymes and the deadenylation machinery (Ciais et al., 2013). To confirm the interaction between POS-1 and MEG-1, we probed the MEG-1::GFP immunoprecipitates with a polyclonal serum against POS-1 (Barbee and Evans, 2006) (Fig. 2B). This experiment confirmed that MEG-1::GFP precipitates contain POS-1, but not the control protein tubulin (Fig. 2B and Fig. S2A,B). POS-1 was not immunoprecipitated by a MEG-3::GFP fusion, further confirming the specificity of the MEG-1-POS-1 interaction (Fig. 2C and Fig. S2C,D). We conclude that MEG-1 exists in a complex that contains P-body components and RNA-binding proteins, including POS-1, a protein predicted to recruit P-body proteins to maternal mRNAs.

### MEG-1 and POS-1 co-localize in P-body-like puncta in P_4_

To examine the distribution of POS-1 and P-body components relative to MEG-1 and P granules, we used antibodies against POS-1 (Barbee and Evans, 2006) and P-body marker DDX6^CGH-1^ (Alessi et al., 2015) and a mNeonGreen::3xFLAG fusion to P-body marker EDC-3 (abbreviated mNG::EDC-3, DeMott et al., 2021). In P_1_ blastomeres, POS-1, DDX6^CGH-1^ and EDC-3 enriched in condensates at the periphery of PGL-3 puncta (Fig. S3A). The POS-1, DDX6^CGH-1^ and EDC-3 condensates overlapped but were not perfectly coincident with MEG-1 (Fig. S3B). In P_4_, MEG-1, POS-1, DDX6^CGH-1^ and EDC-3 appeared to mix more extensively with each other and PGL-3 (Fig. S3A,B). We reasoned that if P-body components associate with MEG-1, they might still form condensates in the absence of P granules. As expected, we found that in *meg-3 meg-4* embryos, which lack P granules, POS-1, DDX6^CGH-1^ and EDC-3 enriched in cytoplasmic puncta most prominently in P_4_, and these co-localized with MEG-1 (Fig. 3A).

**Fig. 3:**
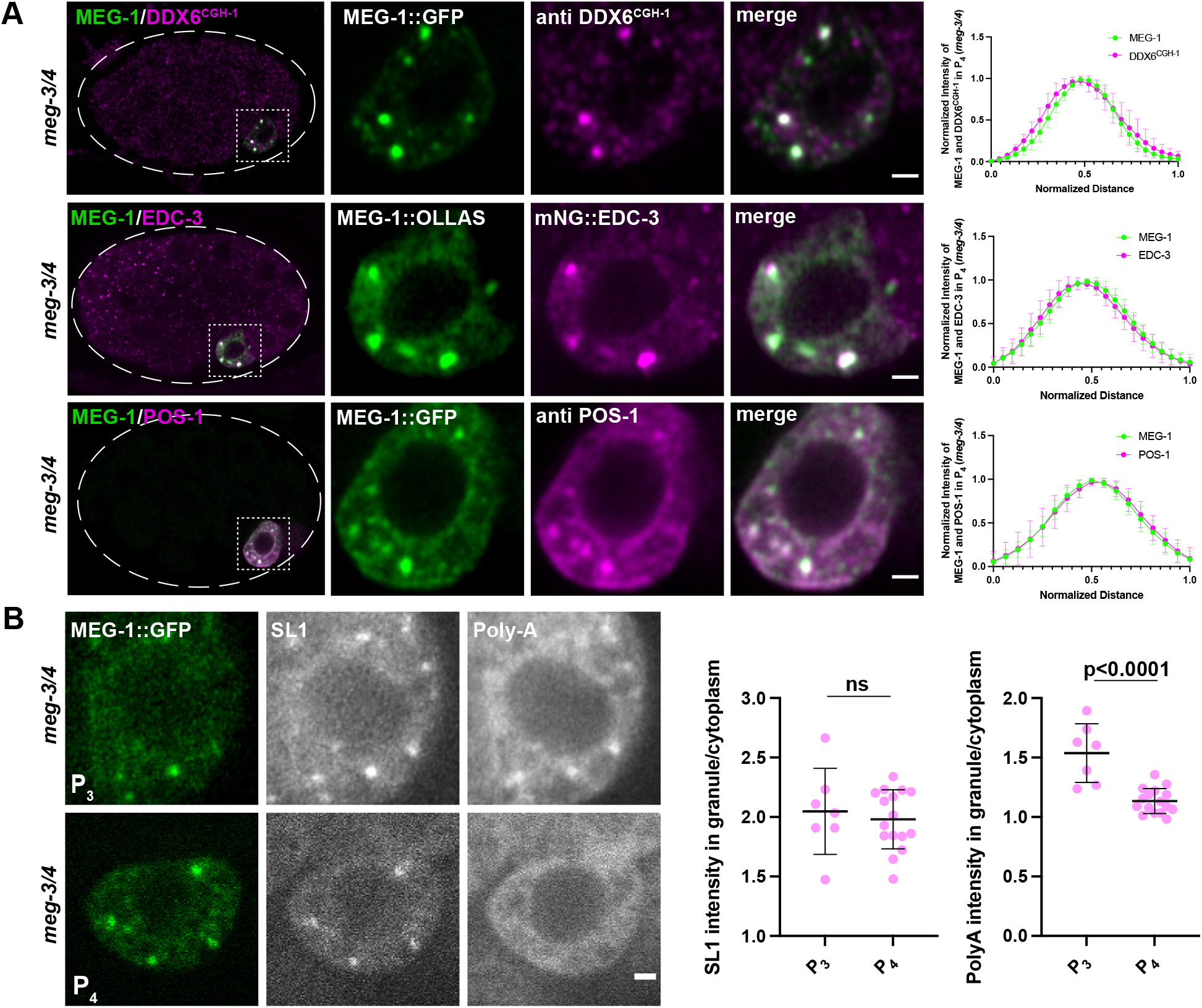
MEG-1 puncta in P_4_ correspond to germline P-bodies. (A) Airyscan photomicrographs of *meg-3 meg-4* embryos expressing MEG-1::GFP and co-stained for GFP and DDX6^CGH-1^, expressing MEG-1::OLLAS and mNG::3xFLAG::EDC-3 and co-stained for OLLAS and FLAG, and expressing MEG-1::GFP and co-stained for GFP and POS-1. Inset shows P_4_ blastomere. Graphs plotting the mean intensities through the center of a granule indicate colocalization. For MEG-1 and DDX6^CGH-1^ n= 7 granules from 2 embryos, for MEG-1 and EDC-3 n=9 granules from 2 embryos, for MEG-1 and POS-1 n=10 granules from 2 embryos. (B) Photomicrographs of *meg-3 meg-4* embryos expressing MEG-1::GFP and probed for SL1 and poly-A. MEG-1 foci enrich SL1 to similar levels in P_3_ and P_4_, but show higher enrichment of poly-A in P_3_ compared to P_4_. The ratio of SL1 or poly-A intensity in MEG-1 granules over cytoplasm in P_3_ (n=7) was compared to P_4_ (n=16). Significance calculated by *t*-test. Quantification for each genotype is from one experiment where several mutant and control animals were processed in parallel. All error bars represent mean ± s.d. All scale bars are 1 μm.

*C. elegans* mRNAs can be detected using an oligo-dT probe that detects poly-adenylated mRNAs and a probe against SL1, the splice leader found on the 5’ end of ~60% of *C. elegans* mRNAs (Seydoux and Fire, 1994). Consistent with enriching maternal mRNAs, P granules are positive for both SL1 and poly-A (Seydoux and Fire, 1994). We reasoned that, since P-bodies are thought to enrich deadenylated mRNAs (Ivanov et al., 2019), P-bodies might be positive for SL1 but not poly-A. P-bodies also assemble in somatic blastomeres, becoming most prominent at the 4-cell stage when degradation of maternal mRNAs begins in somatic lineages (Gallo et al., 2008). Consistent with harboring deadenylated mRNAs, somatic P-bodies marked by EDC-3 showed a high SL1 signal but no poly-A enrichment (compared to the surrounding cytoplasm, Fig. S3C). Similarly, we found that MEG-1::GFP puncta in P_4_ of *meg-3 meg-4* embryos were positive for SL1 but not poly-A (Fig. 3B). Interestingly, MEG-1::GFP puncta in P_3_ were positive for both SL1 and poly-A (Fig. 3B), suggesting that at this stage MEG-1 puncta do not yet correspond to mature P-body-like structures.

Taken together, these observations suggest that, in early P blastomeres, MEG-1 and P-body proteins form overlapping, but not perfectly coincident, assemblies at the periphery of P granules. In P_4_, MEG-1 and P-body components come together into condensates that contain deadenylated mRNAs. We refer to these P_4_-specific condensates as “germline P-bodies” to distinguish these from somatic P-bodies which form in somatic blastomeres and do not contain MEG-1 or POS-1.

### meg-1 and meg-2 are required to maintain DDX6^CGH-1^ and EDC-3 and assemble robust germline P-bodies in P_4_

Unlike P granule proteins, such as PGL-3, which are asymmetrically segregated from the zygote stage (Fig. S1D), DDX6^CGH-1^ and EDC-3 are inherited by all blastomeres during early cleavages. After the 8-cell stage, DDX6^CGH-1^ is turned over in somatic blastomeres (Boag et al., 2005) and remains at high levels only in P_4_ (Fig. S4A-C). EDC-3 is maintained in somatic blastomeres throughout embryogenesis but enriches in P_4_(Fig. S4D-F). In *meg-1 meg-2* mutants, DDX6^CGH-1^ and EDC-3 distributions were unchanged through the 8-cell stage, but DDX6^CGH-1^ was not maintained, and EDC-3 was not enriched, in P_4_ (Fig. S4). In contrast, POS-1, which enriches with germ plasm from the zygote stage (Han et al., 2018), was not affected in *meg-1 meg-2* (Fig. 4A). To quantify these observations, we compared the levels in P_4_ of DDX6^CGH-1^, EDC-3 and POS-1 in *meg-1 meg-2, meg-3 meg-4*, and embryos depleted of all four MEG proteins *[meg-1(vr10) meg-2(RNAi) meg-3(tm4259) meg-4(RNAi)* embryos] (Fig. 4A). DDX6^CGH-1^ and EDC-3 levels were significantly reduced in P_4_ of *meg-1 meg-2* embryos compared to wild-type and in *meg-1 meg-2 meg-3 meg-4* embryos compared to *meg-3 meg-4* embryos (Fig. 4A,B). In contrast, POS-1 levels were not significantly affected in either *meg-1 meg-2* or *meg-3 meg-4* mutants and were reduced only in the quadruple mutant. We conclude that MEG-1/2 are essential to maintain high levels of DDX6^CGH-1^ and EDC-3 in P_4_ and are required redundantly with MEG-3/4 to maintain high levels of POS-1 in P_4_.

**Fig. 4:**
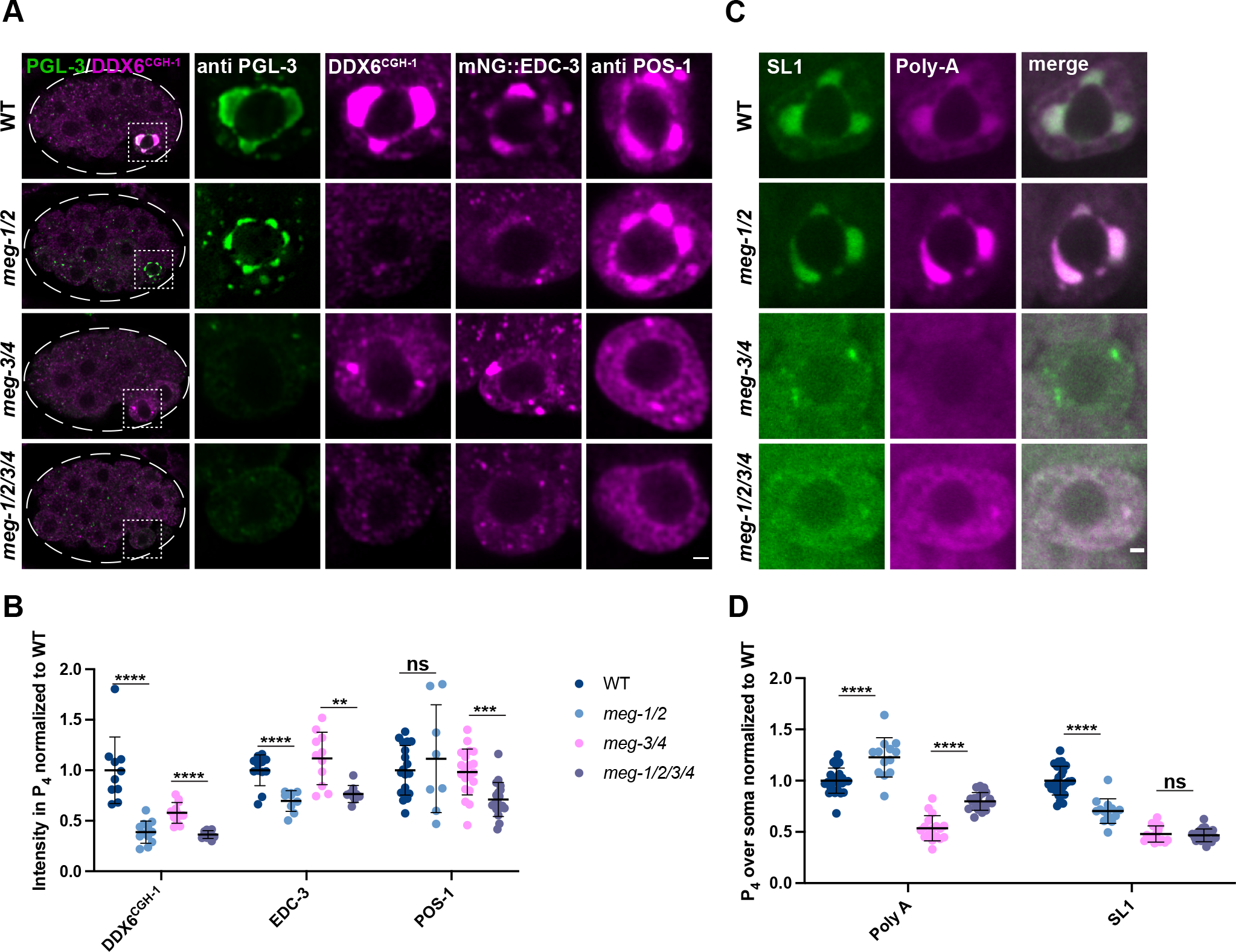
MEG-1/2 are required for maintenance of germline P-bodies in P_4_. (A) Airyscan photomicrographs of embryos of the indicated *meg* genotypes co-stained for PGL-3 and DDX6^CGH-1^ (whole embryo and P_4_ inset), or expressing mNG::3xFLAG::EDC-3 and stained for FLAG, or stained for POS-1. *meg-1 meg-2* are not essential for localization of PGL-3 or POS-1 to P_4_ but are required for maintenance of DDX6^CGH-1^ and EDC-3. (B) Intensity of DDX6^CGH-1^, EDC-3 and POS-1 in P_4_ relative to wild type. Quantification of DDX6^CGH-1^ for each genotype is from one experiment where mutant and control animals were processed in parallel. Wild type n=10; *meg-1/2* n=12; *meg-3/4* n=12; *meg-1/2/3/4* n=10. Quantification of EDC-3 for each genotype is from one experiment where mutant and control animals were processed in parallel. Wild type n=12; *meg-1/2* n=9; *meg-3/4* n=11; *meg-1/2/3/4* n=9. Quantification of POS-1 for *meg-1 meg-2* embryos is from one experiment and from *meg-3 meg-4* and *meg-1 meg-2 meg-3 meg-4* are from two experiments where mutant and control animals were processed in parallel. Wild type n=19; *meg-1/2* n=8; *meg-3/4* n=20; *meg-1/2/3/4* n=19. (C) Photomicrographs of P_4_ in the indicated genotypes probed for SL1 and poly-A. Poly-A levels are increased in *meg-1 meg-2* mutants, despite SL1 levels decreasing or not changing. (D) Quantification of poly-A and SL1 in P_4_ over soma normalized to wild type. Quantification for *meg-1 meg-2* embryos are from two experiments and from *meg-3 meg-4* and *meg-1 meg-2 meg-3 meg-4* are from three experiments where mutant and control animals were processed in parallel. Wild type n=26; *meg-1/2* n=13; *meg-3/4* n=17; *meg-1/2/3/4* n=20. All error bars represent mean ± s.d. \*\*\*\**P*≤0.0001; \*\*\**P*≤0.001; \*\**P*≤0.01; ns= not significant (*t*-test). All scale bars are 1 μm.

The reduction in DDX6^CGH-1^ and EDC-3 levels in P_4_ suggests that germline P-body activity might be compromised in *meg-1 meg-2* mutants. Consistent with this hypothesis, *in situ* hybridization against poly-A and SL1 revealed that poly-A levels were higher in P_4_ of *meg-1 meg-2* embryos compared to wild-type and in *meg-1 meg-2 meg-3 meg-4* embryos compared to *meg-3 meg-4* embryos, despite either a reduction or no significant change in SL1 levels (Fig. 4C,D). We observed SL1+ puncta in P_4_ in 14/17 of *meg-3 meg-4* embryos and in 4/20 *meg-1 meg-2 meg-3 meg-4* embryos (Fig. S5A). The SL1+ puncta did not enrich poly-A over the cytoplasm in *meg-3 meg-4* embryos but did in *meg-1 meg-2 meg-3 meg-4* embryos (Fig. S5A). Together these observations indicate that MEG-1 and MEG-2 are required to maintain robust levels of P-body proteins and robust activation of mRNA deadenylation in P_4_.

### meg-1 meg-2 embryos fail to turnover transcripts targeted for deadenylation by POS-1

To examine directly whether *meg-1 meg-2* mutants exhibit defects in maternal mRNA regulation, we performed RNAseq to compare the transcriptomes of *meg-1 meg-2* mutant embryos to that of wild-type. Two independent RNA-seq libraries were analyzed for each genotype (wild type and *meg-1(vr10) meg-2(RNAi))*. This analysis identified 550 upregulated mRNAs, and 230 downregulated mRNAs, in *meg-1 meg-2* embryos compared to wild-type (±1.5 fold change, *p*<0.05; Fig. 5A, Table S3).

**Fig. 5:**
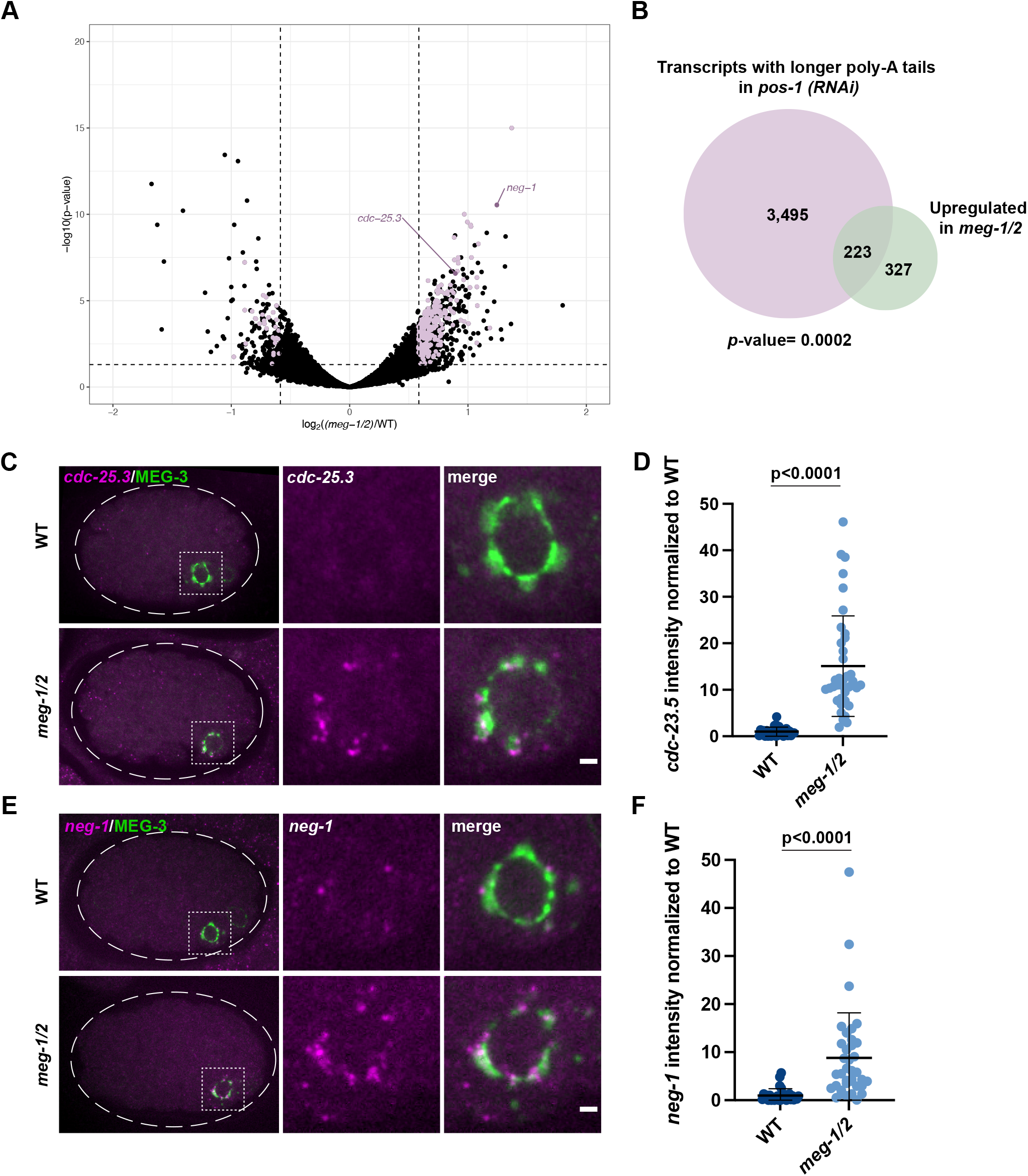
*meg-1/2* are required for the turnover of a subset of POS-1 targets. (A) RNA-seq from two independent experiments comparing *meg-1 meg-2* (RNAi) and wild-type embryos identified 230 downregulated and 550 upregulated genes (± 1.5 fold change). Purple dots correspond to genes significantly down/upregulated in *meg-1 meg-2* embryos that also exhibited longer poly-A tails in *pos-1(RNAi)* embryos (Elewa et al., 2015). (B) 223 genes upregulated in *meg-1 meg-2* embryos overlap with genes whose poly-A tails were extended in *pos-1*(RNAi) embryos P=0.0002 (Fisher’s exact test, see methods). (C) and (E) Photomicrographs of *cdc-25.3* and *neg-1* smFISH in embryos expressing the P granule marker MEG-3::GFP. Inset shows P_4_. *cdc-25.3* and *neg-1* are turned over less efficiently in *meg-1 meg-2* P_4_ blastomeres. Scale bars are 1 μm. (D) and (F) Intensity of *cdc-25.3* and *neg-1* in P_4_ normalized to wild type. *In situs* for *cdc-25.3* and *neg-1* were done in the same embryos in two independent experiments where mutant and control animals were processed in parallel. Wild type n=29; *meg-1/2* n=38. Error bars represent mean ± s.d. A *t*-test was used to make comparisons between genotypes.

Elewa et al., 2015 identified 3,726 transcripts that display longer poly-A tails in *pos-1*(RNAi) embryos compared to wild-type (“deadenylated POS-1 targets”), of which 3,718 were detected in our RNA-seq. 40% of genes (223/550) upregulated in *meg-1 meg-2* embryos were among these deadenylated POS-1 targets (Fig. 5B, Table S3). Assuming a total pool of 11,121 transcripts that can be detected by these analyses in early embryos (see Methods), we found this overlap to be significant (Fisher’s exact test, *p*-value=0.0002). In comparison, the overlap between transcripts downregulated in *meg-1 meg-2* embryos and deadenylated POS-1 targets (30/3,718 transcripts; *p*-value = 1) or adenylated POS-1 targets (transcripts with shorter poly-A tails in pos-1(RNAi); 17/1,307; *p*-value=0.99) was not significant (but see next section). We conclude that MEG-1 and MEG-2 contribute to the turn-over of a subset of maternal mRNAs also targeted by POS-1 for deadenylation.

*neg-1* and *cdc-25.3* are two transcripts among the 223 potential targets shared between POS-1 and MEG-1. *neg-1* and *cdc-25.3* are maternally-deposited and turned over in all lineages by the 28-cell stage (Fig. S6A,B; Tintori et al., 2016; Elewa et al., 2015). In *meg-1 meg-2* embryos, but not in *meg-3 meg-4* embryos, *neg-1* and *cdc-25.3* transcripts were still detected in P_4_ in the 28-cell stage (Fig. 5C-F and Fig. S6C,D). These observations confirm that *meg-1/2* activity is required for the efficient turnover of a subset of POS-1-regulated transcripts.

### meg-1 meg-2 embryos fail to express efficiently transcripts activated by POS-1 for translation in P_4_

In addition to promoting deadenylation of a subset of maternal transcripts, POS-1 is also required to extend the poly-A tail of a different group of maternal transcripts that are translationally activated in embryos, including *nos-2, Y51F10.2 and xnd-1* (Elewa et al., 2015). These transcripts code for factors required for germ cell fate and are translationally repressed in the P_0_, P_1_, P_2_ and P_3_ blastomeres and translationally activated in P_4_(Lee et al. 2020; Mainpal et al., 2015). Translational activation of *nos-2* and *Y51F10.2* has been confirmed to require POS-1 (D’Agostino et al., 2006; Jadhav et al., 2008; Lee et al., 2020).

We used *in situ* hybridization and immunofluorescence to examine transcript and protein levels in P_4_ of wild type, *meg-1 meg-2, meg-3 meg-4* and *meg1 meg-2 meg-3 meg-4* embryos (Fig. 6). We found that for all three transcripts, RNA levels were lowest in the *meg-3 meg-4* mutants, consistent with a dependence on P granules for enrichment in P_4_. RNA levels were also reduced in *meg-1 meg-2* mutants compared to wild-type, suggesting that MEG-1/2 also contribute to RNA enrichment either directly or indirectly through an effect on P granule segregation, since P granules are also inefficiently segregated in these mutants (Fig. S1D). Adjusting for RNA levels, we found that protein output was reduced in *meg-1 meg-2* and elevated in *meg-3 meg-4* compared to wild-type (Fig. 6). These differences did not correlate with POS-1 protein levels in P_4_, which were similar in these mutants (Fig. 4 A,B). Consistent with *meg-1 meg-2* and *meg-3 meg-4* acting in parallel, protein levels were lowest in embryos depleted of all four *megs* compared to either double combination. Together, these observations suggest that *meg-1 meg-2* and *meg-3 meg-4* contribute independently to expression of maternal transcripts in P_4_, with MEG-3/4 acting primarily by boosting RNA levels and MEG-1/2 primarily by boosting protein output.

**Fig. 6:**
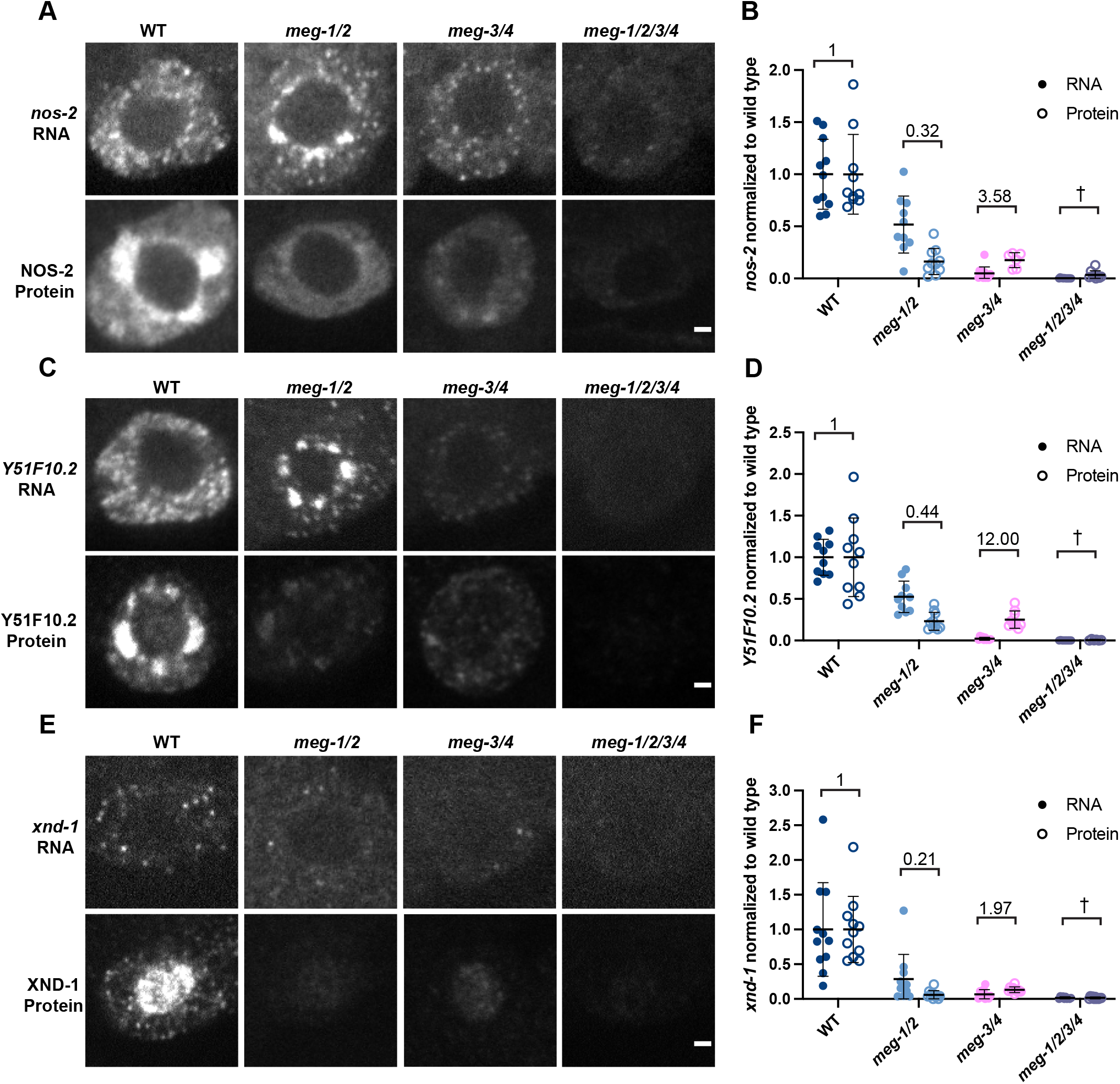
*meg-1/2* are required for efficient translation of maternal mRNAs coding for germ cell fate determinants. (A), (C) and (E) Photomicrographs of P_4_ in embryos of the indicated genotypes comparing *nos-2, Y51F10.2*, and *xnd-1* RNA and protein levels. In all cases, the RNA is partially reduced in *meg-1 meg-2* mutants, and dramatically reduced in *meg-3 meg-4* and *meg-1 meg-2 meg-3 meg-4*. In contrast, the protein levels of *meg-1 meg-2* and *meg-3 meg-4* are similar. Scale bars are 1 μm. (B), (D) and (F) Intensity of RNA and protein, normalized to wild type. The ratio of protein to RNA levels in each genotype is indicated. In *meg-1 meg-2*, the ratio is decreased, while in *meg-3 meg-4* it is increased. † Due to the very low levels of RNA present in *meg-1 meg-2 meg-3 meg-4* embryos we were unable to calculate the protein/RNA ratio. Quantification for each genotype is from one experiment where mutant and control animals were processed in parallel. For *nos-2* RNA: wild type n=11, *meg-1/2* n=10, *meg-3/4* n=12, *meg-1/2/3/4* n=12. For NOS-2 protein: wild type n=10, *meg-1/2* n=10, *meg-3/4* n=6, *meg-1/2/3/4* n=9. For *Y51F10.2* RNA: wild type n=10, *meg-1/2* n=10, *meg-3/4* n=10, *meg-1/2/3/4* n=9. For *Y51F10.2* protein: wild type n=10, *meg-1/2 n*=10, *meg-3/4* n=9, *meg-1/2/3/4* n=6. For *xnd-1* RNA: wild type n=11, *meg-1/2* n=11, *meg-3/4* n=10, *meg-1/2/3/4* n=10. For XND-1 protein: wild type n=11, *meg-1/2* n=11, *meg-3/4* n=10, *meg-1/2/3/4* n=11. Error bars represent mean ± s.d.

In wild type, *nos-2* and *Y51F10.2* RNAs enrich in P granules through P_3_ and become cytoplasmic in P_4_ coincident with translational activation (Lee et al., 2020). *[xnd-1* is a much less abundant transcript which precluded us from evaluating its partitioning between P granules and the cytoplasm (Fig. 6E)]. Consistent with reduced translational activation in P_4_, we observed that *nos-2* and *Y51F10.2* remained enriched in a perinuclear pattern in *meg-1 meg-2* embryos, as also observed in *pos-1* embryos (Lee et al., 2020) (Fig. 6A,C, Fig. S7). As mentioned above, *nos-2* and *Y51F10.2* exhibited a higher protein output in P_4_ in *meg-3 meg-4* embryos compared to wild-type and *meg-1 meg-2* embryos (Fig. 6B and 6D), suggesting that assembly into P granules dampens translational activation. We could not determine translational output in *meg-1 meg-2 meg-3 meg-4* due to the extremely low levels of RNA in P_4_ in these mutants. We conclude that *meg-1 meg-2* are required for maximal translation activation of POS-1 targets in P_4_, which is antagonized by *meg-3 meg-4*.

### Primordial germ cells adopt a muscle precursor-like cell fate in meg-1 meg-2 mutants

In *pos-1* mutants, P_4_ descendants develop as muscle precursor cells that express the myoD homolog *hlh-1* (Tabara et al., 1999). To determine whether a similar cell fate transformation occurs in *meg-1 meg-2* mutants, we examined the expression of *hlh-1* and the PGC zygotic transcript *xnd-1* (Mainpal et al., 2015) by *in situ* hybridization using a P granule marker to identify P_4_ descendants. We observed *hlh-1* transcripts in P_4_ descendants in 21/23 bean to comma stage *meg-1 meg-2* embryos examined, compared to 0/21 wild-type embryos examined (Fig. 7A). In contrast, we failed to observe robust expression of *xnd-1* in 16/24 *meg-1 meg-2* embryos (Fig. 7B).

**Fig.7:**
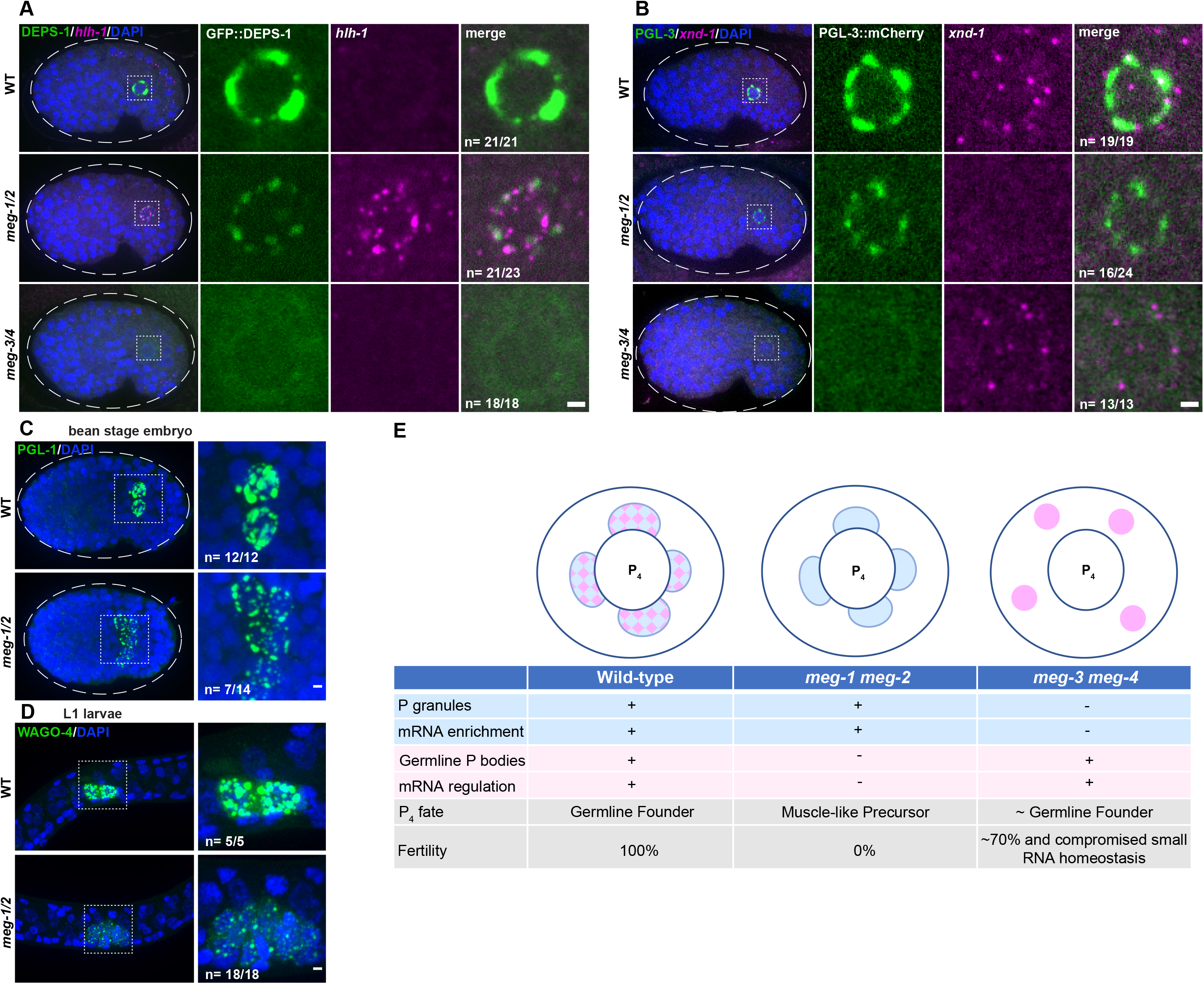
Primordial germ cells exhibit somatic-like characteristics in *meg-1 meg-2* mutants. (A) Photomicrographs of bean stage embryos of the indicated genotypes expressing DEPS-1::GFP and probed for *hlh-1* RNA. Inset depicts a primordial germ cell. Embryos were scored from one independent experiment where mutant and control animals were processed in parallel. 21/21 wild-type and 18/18 *meg-3 meg-4* bean to comma stage embryos did not express *hlh-1*. 21/23 *meg-1 meg-2* did express *hlh-1*. (B) Photomicrographs of bean stage embryos of the indicated genotypes expressing PGL-3::mCherry and probed for *xnd-1* RNA (which is transcribed in PGCs at this stage). Inset depicts a primordial germ cell. Embryos were scored from two independent experiments for *meg-1 meg-2* and one experiment for *meg-3 meg-4* where mutant and control animals were processed in parallel. 19/19 wild-type and 13/13 *meg-3 meg-4* bean stage embryos expressed *xnd-1*. 16/24 *meg-1 meg-2* embryos did not express *xnd-1*. (C) Maximum projections of bean stage embryos of the indicated genotypes stained for PGL-1. Inset shows the primordial germ cells. Embryos were scored from one experiment where mutant and control animals were processed in parallel. 12/12 wild-type embryos had two PGL-1 positive cells and 7/14 *meg-1 meg-2* embryos had more than two PGL-1 positive cells. (D) Maximum projections of germ cells from unfed L1 larvae expressing the germ granule marker 3xFLAG::GFP::WAGO-4. Embryos were scored from one experiment where mutant and control animals were processed in parallel. 5/5 wild-type embryos had two WAGO-4 positive cells and 18/18 *meg-1 meg-2* embryos had more than two WAGO-4 positive cells. All scale bars are 1 μm. (E) Working model: Cartoon and table summarizing P_4_ phenotypes based on this study and on Wang et al., 2014 and Ouyang et al., 2019. P granules are depicted in blue, germline P-body in pink, and their merge in a checkered pattern. Note that P granule and germline P-body proteins also exist in a more dilute state in the cytoplasm. See text for additional details.

In wild type, the daughters of P_4_ (Z2 and Z3) remain non-proliferative during embryogenesis and only divide in L1 larvae after the onset of feeding. In *meg-1 meg-2* mutants, we observed more than two P granule-positive cells in 50% of bean-to-comma cell stage embryos (Fig. 7C) and in 100% of non-fed L1 larvae stage (Fig. 7D). The extra P granule-positive cells were not due to miss-segregation of P granules to the D blastomere (Fig. S8A), were first detected around the 35-45 cell stage (Fig. S8B) and did not express muscle myosin (Fig. S8C). We conclude that primordial germ cells are partially transformed to muscle precursor-like fate in *meg-1 meg-2* mutants.

The *meg-1 meg-2* phenotype contrasts with that of *meg-3 meg-4* embryos where Z2 and Z3 express *xnd-1*, do not express *hlh-1* and do not proliferate prematurely despite the absence of maternal P granules (Fig. 7A,B; Wang et al., 2014). ~70% of *meg-3 meg-4* mutants are fertile, in contrast to *meg-1 meg-2* mutants which are 100% sterile (Leacock and Reinke et al., 2008; Wang et al., 2014).

## Discussion

In this study, we demonstrate that the germ plasm of *C. elegans* contains two condensate types, P granules and germline P-bodies. Each rely on a different pair of intrinsically-disordered proteins for efficient accumulation in the germline founder cell P_4_: P granules depend on MEG-3 and MEG-4 and germline P-bodies depend on MEG-1 and MEG-2. We used these distinct genetic requirements to distinguish the contribution of each condensate to germ cell fate (Fig. 7E). P granules enrich regulators of small RNA homeostasis (Ouyang et al., 2019) and maternal mRNAs but are not required for maternal mRNA regulation (Lee et al, 2020 and this study). mRNA regulation depends on “germline P-bodies”, which promote the translation of mRNAs coding for germline determinants and the turn-over of mRNAs coding for somatic determinants. We propose that the germ cell fate-specifying “germ granules” of *C. elegans* are assemblies of at least two distinct condensates, P granules and germline P-bodies, which enrich and regulate, respectively, maternal mRNAs in the germline founder cells.

### Germline P-bodies and P granules are two types of condensates that require MEG proteins for stabilization in the embryonic germ lineage

P granules were the first characterized condensates in the *C. elegans* germ plasm (Strome and Wood, 1982). P granules consist of a dense liquid core, assembled by PGL proteins, surrounded by interfacial nanoscale, RNA-rich solid clusters assembled by intrinsically-disordered proteins MEG-3 and MEG-4 (Folkmann and Putnam et al., 2021). In this study, we describe a second condensate type, germline P-bodies, that contains regulators of mRNA adenylation and decapping, the RNA-binding protein POS-1, and MEG-1 and MEG-2, two intrinsically-disordered proteins related to MEG-3 and MEG-4. Germline P-body components assemble in complex patterns around P granules in early P blastomeres and merge with each other and P granules in P_4_. In embryos lacking P granules *(meg-3 meg-4* mutants), germline P-bodies can be visualized in P_4_ as discrete SL1+ poly-A-cytoplasmic puncta that are also positive for MEG-1, POS-1 and the canonical P-body markers DDX6^CGH-1^ and EDC-3. In the absence of *meg-1 meg-2*, DDX6^CGH-1^ and EDC-3 levels are reduced and maternal mRNA regulation fails, despite normal P granule assembly and POS-1 levels (Fig. 7E).

How MEG-1/2 stabilize germline P-body components remains unclear. Unlike MEG-3/4 which are required for the asymmetric segregation of P granules from the zygote stage onward, MEG-1/2 do not appear to affect the distribution of germline P-body components until after the 8-cell stage. P-body components (DDX6^CGH-1^ and EDC-3) are initially segregated to all cells and coalesce into puncta in somatic cells coincident with the onset of maternal mRNA degradation (Gallo et al., 2008). MEG-1/2 do not affect P-body assembly in somatic cells but are required for stabilization of DDX6^CGH-1^ and EDC-3 specifically in P_4_, at the embryonic stage when DDX6^CGH-1^ is rapidly cleared from somatic lineages. In *Drosophila* embryos, the DDX6/4-ET-like complex (ME31B/Cup) is targeted for degradation by CTLH, an E3 ubiquitin ligase, and Marie Kondo, an E2 conjugating enzyme (Cao et al., 2020; Zavortink et al., 2020). It will be interesting to determine whether homologs of these factors promote DDX6^CGH-1^ turnover in *C. elegans* and how MEG-1/2 might oppose these activities in P_4_.

In contrast to somatic blastomeres which activate zygotic transcription by the 4-cell stage, P blastomeres remain transcriptionally silent until the birth of the daughters of P_4_, the primordial germ cells Z2 and Z3 (100-cell stage). We suggest that MEG-enhanced condensation of P granules and germline P-bodies serves as a mechanism to concentrate maternally-provided mRNAs and their regulators in germ plasm to ensure that P_4_ inherits sufficient machinery to initiate the maternal-to-zygotic transition. The MEG-1/2 and MEG-3/4 paralog pairs appear to have diverged such that MEG-1/2 interact preferentially with P-body components and MEG-3/4 interact preferentially with P granule components. MEG-3/4, but not MEG-1/2, contain an HMG-like domain essential for MEG-3/4 clusters to associate with the surface of PGL condensates (Schmidt et al., 2021). MEG-3/4 stabilize PGL condensates by lowering their surface tension (Folkmann and Putnam et al., 2021); it remains to be determined whether MEG-1/2 function similarly or by another mechanism.

### Germline P-body proteins control maternal mRNA regulation in the germline founder cell P_4_

The birth of the P_4_ blastomere appears to coincide with a major transition in maternal mRNA regulation in the P lineage as evidenced by 1) coalescence of germline P-bodies containing deadenylated mRNAs, 2) degradation of transcripts coding for somatic factors, and 3) translation of transcripts coding for germ cell fate determinants. We suggest that regulators of mRNA adenylation and decapping that enrich in P-bodies drive this transition in P_4_ by targeting maternal mRNAs for de-adenylation/degradation or adenylation/translation, depending on the combination of RNA-binding proteins, including POS-1, bound to 3’ UTRs. The poly-A polymerase subunits GLD-2 and GLD-3 are enriched in MEG-1 immunoprecipitates and have been reported to enrich in granules in germ plasm (Wang et al., 2002; Eckmann et al., 2002). It will be interesting to determine whether GLD-2/3 also localize to germline P-bodies and are responsible for the translational activation of transcripts like *nos-2, Y51F10.2* and *xnd-1*.

The birth of P_4_ also coincides with the apparent mixing of germline P-bodies and P granules and the release of *nos-2* and *Y51F10.2* mRNAs from P granules coincident with their translational activation. This is also the stage where Z granules and SIMR-1 foci appear to de-mix from P granules to form the multi-condensate nuage characteristic of pre-gametic germ cells (Wan et al., 2018; Uebel et al., 2021). These observations suggest a dramatic switch in the material properties of condensates in the transition from P_3_ to P_4_. We do not know whether these changes arise as a cause, or consequence, of the changes in mRNA regulation that also occur at this stage. In principle, segregation of maternal mRNAs and their regulators into distinct condensates that eventually merge in P_4_ could be used as a physical mechanism to control RNA-protein interactions. Alternatively, changes in condensation patterns could derive from changes in the composition and solubility of complexes dispersed throughout the cytoplasm. We favor the latter since 1) RNAs and proteins enriched in P granules and P-bodies are also found dispersed throughout the cytoplasm and 2) failure to assemble P granules does not prevent timely translational regulation of mRNAs enriched in P granules. We suggest that the complex condensation patterns of germline P-body components in early P blastomeres, and apparent “mixing” with P granules in P_4_, are mesoscale manifestations of molecular-scale rearrangements that occur throughout the cytoplasm and eventually culminate in the targeting of the P-body machinery onto maternal mRNAs in P_4_. What regulates these changes during developmental time remains a mystery. The significance of the close association of germline P-bodies with P granules is also unclear and may reflect the fact that the two condensate types likely share some components such as POS-1, which depends on both MEG-1/2 and MEG-3/4 for maximal segregation to P_4_ (Fig. 4B).

### A conserved role for P-body proteins in specifying germ cell fate

In *meg-1 meg-2* mutants, P_4_ descendants divide precociously, fail to activate the transcription of the germ cell transcript *xnd-1* and activate instead the transcription of the muscle transcription factor MyoD^HLH-1^. These observations suggest a transformation to a muscle precursor fate, such as that normally adopted by the sister of P_4_, the somatic blastomere D. This fate transformation occurs despite maintenance of P granules in Z2 and Z3 and their descendants, confirming that P granules are neither sufficient nor required to specify germ cell fate in primordial germ cells (Gallo et al., 2010; Strome et al., 1995). A similar P_4_ → D fate transformation was reported for *pos-1* mutants (Tabara et al., 1999). The apparent P_4_→ D fate transformation is likely incomplete as Z2 and Z3 descendants do not express muscle myosin, remain in their normal central position in embryos and first-stage larvae, and stall proliferation during the first larval stage. *meg-1 meg-2* fail to efficiently translate NOS-2 and Y51F10.2, two proteins implicated, respectively, in mRNA and protein turnover (Subramaniam and Seydoux, 1999; Kipreos, 2005). We showed previously that the sterility of embryos lacking Nanos could be rescued by reducing the activity of maternal LIN-15B, a soma-promoting transcription factor expressed in oocytes (Lee et al., 2017). Similarly, the germ cell proliferation defect of *meg-1 meg-2* larvae could be rescued partially by reducing *gld-1* activity (Kapelle and Reinke, 2011), an RNA-binding protein required for oocyte development and expressed in early P blastomeres (Francis et al., 1995; Jones et al., 1996). Together these observations suggest that a key step to specify P_4_ as the germline founder cell is to program germline P-bodies to eliminate maternal factors that function during oogenesis.

The germline P-bodies we describe here share several features with the recently described “founder granules” in *Drosophila* germ plasm. Founder granules contain DDX6^ME31B^ the decapping factor DCP1 and Oskar mRNA, which although required for germ plasm assembly in oocytes, must be degraded in embryos for proper germline development (Eichler et al., 2020). DDX6^ME31B^ has been proposed to enrich in germ plasm independently of the canonical Oskar polar granule assembly pathway (McCambridge et al., 2020), as we demonstrate here for germline P-bodies, which assemble independently of P granules. Founder granules, however, have not yet been implicated in the translational activation of Nanos and other mRNAs enriched in polar granules, as we suggest here for germline P-bodies.

A role for P-bodies in early germ cell development has also been suggested by studies in mice. The mammalian Nanos homolog NANOS2 localizes to P-bodies, interacts with the CCR4-NOT1 deadenylation complex, and promotes mRNA degradation and the male germ cell fate program in mice (Suzuki et al., 2010; Shimada et al., 2019; Wright et al., 2021). DDX6/Me31B RNA helicases have also been implicated in the differentiation of various stem cell populations in human, mouse, and *Drosophila* (Di Stefano et al., 2019; Nicklas et al., 2015; Jensen et al., 2021). Together these studies suggest a conserved role for P-bodies as essential regulators of cell fate transitions in progenitors of the germline and beyond.

### Limitations of the study

We inferred a requirement for P-body activity in embryonic germ cells through our analyses of *meg-1 meg-2* mutants which fail to stabilize germline P-bodies and regulate maternal mRNAs in P_4_. We did not test directly, however, for a requirement for P-body enzymatic activity, as mutants in key P-body proteins arrest development before the birth of P_4_. For example, RNAi reduction of the scaffold C-NOT1^NTL-1^ leads to early embryonic division defects, presumably because P-bodies also regulate the fate of mRNAs in somatic blastomeres (Gallo et al., 2008). The helicase DDX6^CGH-1^ stabilizes translationally repressed mRNAs during oogenesis and is essential for the production of mature oocytes that support normal embryogenesis (Boag et al., 2008; Noble et al., 2008). A DDX6^CGH-1^ temperature-sensitive mutant is available (Scheckel et al., 2012), which could potentially allow us to bypass an earlier requirement for DDX6^CGH-1^, but initial experiments proved inconclusive. Although we demonstrate that MEG-1 can be immunoprecipitated from lysates in a complex with POS-1 and a subset of P-body proteins, we have not investigated whether MEG-1 binds directly to these proteins or interacts indirectly by binding RNA for example. We also do not address whether MEG-1/2 or germline P-bodies are merely required (permissive) or are sufficient (instructive) to specify germ cell fate. MEG-1/2 enrich preferentially into P blastomeres from the zygote-stage onward; mutations that prevent this localization may help determine whether MEG-1/2 play a permissive or instructive role in germ cell fate specification.

## Acknowledgments

We thank John Kim and Amelia Alessi, Tom Evans, and Judith Yanowitz for the CGH-1, POS-1, and XND-1 antibodies; Dominique Rasoloson and Helen Schmidt for the MEG-1::GFP and MEG-1::OLLAS alleles; Tu Lu for the strains JH3472, JH3410, JH3404 and JH3352; the⍰Johns Hopkins Microscope Facility (S10OD023548) for microscopy support; the Johns Hopkins University School of Medicine Genetic Resources Core Facility for sequencing support, and the JHMI Mass Spectrometry and Proteomics Facility for mass-spec support. We also thank the Seydoux lab and the Baltimore Worm Club for their insights during this project. Some strains were provided by the CGC, which is funded by NIH Office of Research Infrastructure Programs (P40 OD010440). Funding was provided by the National Institutes of Health (GS: R37HD037047; MC: T32 GM007445) and the National Science Foundation (MC: DGE-1746891). GS is an investigator of the Howard Hughes Medical Institute.

## Data availability

Sequencing data has been deposited onto the Gene Expression Omnibus (GEO) and can be found using the following accession numbers:

#########

Mass spectrometry data has been deposited to the MassIVE repository and can be found with the identifier #########.

## Methods

### Worm handling, RNAi, sterility counts

*C. elegans* were cultured according to standard methods (Brenner, 1974). Strains used in this study are listed in Table S4. RNAi knockdown experiments were performed by feeding on HT115 bacteria (Timmons and Fire, 1998). The empty pL4440 vector was used as a negative control. Bacteria were grown at 37°C in LB + ampicillin (100 μg/mL) media for 5 hours, induced with 5 mM IPTG for 30 minutes, plated on NNGM (nematode nutritional growth media) + ampicillin (100 μg/mL) + IPTG (1 mM) plates, and grown overnight at room temperature. L4 hermaphrodites were put onto RNAi plates and fed overnight at 25°C, and then shifted back to 20°C for at least one hour before proceeding with further experiments. Effectiveness of knocking down *meg* genes was verified by scoring the sterility of adult progeny of the worms exposed to RNAi.

To culture larger numbers of worms, worm cultures were started from synchronized L1s (hatched from embryos incubated in M9 overnight) onto NA22 or RNAi bacteria containing plates and grown to gravid adults at 20°C. Early embryos were harvested from gravid adults.

To measure maternal-effect sterility of the *meg-1 meg-2(ax4532)* strain, 20 gravid adults from a mixed heterozygous population were singled out onto individual OP50 plates. Worms were allowed to lay eggs for 5 hours, then removed and genotyped by PCR. Adult progeny were scored for empty uteri (white sterile phenotype).

### CRISPR genome editing

Genome editing was performed using CRISPR/Cas9 as described in Paix et al., 2017. The *meg-1 meg-2* open reading frame was deleted with two guide RNAs targeting the following sequences: 1. tgagcggcgatggataatcg and 2. agtcaaaattagttgctggg. Deletion of *meg-1 meg-2* was confirmed by Sanger sequencing. This strain (JH3875) is maintained as a heterozygote because the homozygous *meg-1 meg-2* deletion is 100% maternal effect sterile.

### RNA extraction and preparation of mRNA-seq library

For each replicate, 26,000 synchronized L1 worms were plated on HT115 bacteria transformed with either L4440 (control) or *meg-2* RNAi and grown at 20°C until the young adult stage. Adult worms were collected by filtering and the embryos were harvested by bleaching. Embryo pellets were flash frozen in liquid nitrogen. RNA was extracted with TRIzol reagent and chloroform. RNA was then concentrated and purified using Zymo’s RNA Clean & Concentrator kit.

For mRNA-seq library preparation, 1 μg of total RNA was treated with Ribo-Zero Gold rRNA Removal Kit. A 1:100 dilution of ERCC RNA Spike-in Mix was added. Libraries were prepared using the TruSeq stranded total RNA library Prep Kit with 12 cycles of PCR amplification. All sequencing was performed using the Illumina HiSeq2500 at the Johns Hopkins University School of Medicine Genetic Resources Core Facility.

### mRNA-sequencing analysis

Sequencing reads were aligned to the UCSC ce10 *C. elegans* reference genome using HISAT2 (Kim et al., 2015). Reads aligning to genetic features were then counted using HTSeq-count (Anders et al., 2015) and analyzed for differential expression analysis using DESeq2 (Love et al., 2014). Genes differentially expressed in wild-type vs *meg-1 meg-2* embryos are listed in Table S3.

### Immunoprecipitation

For each replicate for mass spec analysis, 1×10^6^ synchronized L1 worms were grown on NA22 bacteria at 20°C until the young adult stage. For IPs to compare MEG-1::GFP and MEG-3::GFP by western blotting, 4x as many MEG-3::GFP embryos were collected as MEG-1::GFP embryos, because MEG-1 is ~4x more abundant than MEG-3 (Saha et al., 2016). Adult worms were collected by filtering and the embryos were harvested by bleaching. Embryos were washed and flash frozen in IP buffer (300 mM KCl, 50 mM HEPES pH 7.4, 1 mM EGTA, 1 mM MgCl_2_, 1% glycerol, 0.1% NP-40) with 2x freshly prepared protease inhibitor mix #1 and mix #2 (100x protease inhibitor mix #1 contained 3 mg/mL antipain, 5 mg/mL leupeptin, 10 mg/mL benzamidine, 25 mg/mL AEBSF, and 1 mg/mL phosphoramidon diluted in PBS. 100x protease inhibitor mix #2 contained 5 mg/mL aprotinin, 4 mM bestatin, 1 mg/mL E64 and 1 mg/mL trypsin inhibitor diluted in water). Thawed embryos were sonicated on ice with a Branson Digital Sonifier SFX 250 with a microtip (15s on, 45s off, 15% power, 6 minutes total on time or until embryos were completely lysed) and cleared by centrifugation at 4°C for 30 minutes at 20,817 RCF.

For the IP, 150 μl of anti-GFP nanobody conjugated to magnetic beads (Chromotek; cat# gtma-10) were incubated with the lysates at 4°C for 90 minutes. The unbound fraction was removed and the beads were washed five times with ice cold IP buffer. The bound fraction was eluted by boiling the beads in 1% SDS with 50mM Tris-HCL pH 7.4 for 5 minutes.

### Western blotting

1 M DTT and NuPAGE LDS sample buffer(4x) were added to lysates to a final concentration of 200 mM DTT and 1x NuPAGE LDS sample buffer. Samples were boiled for 5 minutes and run on 4-12% Bis-Tris gels in MES buffer. Samples were transferred to a PVDF membrane. Membranes were blocked in PBS with 0.1% Tween 20 and 5% non-fat dry milk (PBST + 5% milk). Membranes were incubated in primary antibodies diluted in PBST + 5% milk overnight at 4°C. Membranes were washed three times for 10 minutes in PBST and then incubated with secondary antibodies diluted in PBST + 5% milk at room temperature for 1 hour. Membranes were washed again three times for 10 minutes in PBST and visualized with Pierce ECL Western Blotting Substrate (Thermo; cat# 32106) or SuperSignal West Femto Maximum Sensitivity Substrate (Thermo; cat# 34095) and the KwikQuantTM Imager (Kindle Biosciences).

Primary antibodies and concentrations used: mouse anti-GFP Living Colors (JL-8) (Takara Biosciences; cat# 632381) 1:500 dilution. Mouse anti-α-Tubulin (Sigma; cat# T6199) 1:1,000 dilution. Rabbit anti-POS-1 (a gift from Tom Evans) 1:500 dilution.

### Mass spectrometry

Mass spectrometry was performed by the JHMI Mass Spectrometry and Proteomics Facility. Samples were reduced with DTT, alkylated with iodoacetamide, TCA/acetone precipitated, and in solution digested with trypsin. Samples were analyzed by LC-MS-MS on Q-Exactive Plus (Thermo) in FTFT at resolution 140K/35K with total 120 minute gradient.

### Mass spec data analysis

Raw data were processed and analyzed using MaxQuant (2.0.3.0) software (Tyanova et al., 2016a). Default settings were used except that ‘Match between runs’ was turned on. Search parameters were as follows: Cysteine carbamidomethyl was included as a fixed modification, and variable modifications included oxidation of methionine, protein N-terminal acetylation, deamidation of glutamine and asparagine, and phosphorylation of serine, threonine and tyrosine, and the maximum number of modifications per peptide was set to 4. Trypsin was used as the digestion enzyme, a maximum of two missed cleavages were allowed, and the minimal peptide length was set to seven amino acids. Database search was performed against Uniprot *C. elegans* database (UP000001940_6239.fasta). False discovery rate (FDR) was set to 1% at peptide spectrum match (PSM) and protein level. Minimum peptide count required for protein quantification was set to two. Protein groups were further analyzed using Perseus (Tyanova et al., 2016b). Common contaminants, reverse proteins and proteins only identified by site were filtered out. LFQ values were log_2_ transformed. Two-sample *t-tests* were performed.

### Immunostaining

Embryos were extruded from adult animals and subjected to freeze-crack on 0.01% poly-lysine coated slides followed by fixation in −20°C methanol ≥ 15 minutes. Slides were blocked in PBS with 0.1% Tween 20 and 0.1% BSA (PBST + BSA) for 1 hour. Slides were incubated in primary antibodies diluted in PBST + BSA at 4°C in a humidity chamber overnight. Slides were washed three times in PBST for 5 minutes and then incubated in secondary antibodies diluted in PBST + BSA for 1 hour at room temperature. Slides were washed again three times in PBST for 5 minutes, then two quick washes in PBS. Samples were mounted in ProLong Glass Antifade mountant and cured overnight. When co-staining with OLLAS antibody, the OLLAS primary and secondary were applied first to avoid cross reactions.

Primary antibodies and concentrations used: Mouse anti-FLAG M2 (Sigma; cat# F1804) 1:500. Rat anti-OLLAS L2 (Novus; cat# 06713) 1:50. Rabbit anti-CGH-1 (a gift from John Kim; Alessi et al., 2015) 1:1,000. Rabbit anti-POS-1 (a gift from Tom Evans; Barbee and Evans, 2006) 1:100. Guinea pig anti-XND-1 (a gift from Judith Yanowitz; Wagner et al., 2010) 1:2,000. Mouse anti-PGL-3 KT3 (DSHB) 1:100. Mouse anti-PGL-1 OIC1D4 (DSHB) 1:10. Mouse anti-UNC-54 mAB 5-8 (DSHB) 1:10. Anti-GFP nanobody conjugated to Alexa Fluor 488 (Chromotek; cat# gb2AF488-10) 1:500. Antibody staining in this manuscript was consistent with that of previously published works.

### Single molecule fluorescence in situ hybridization (smFISH)

smFISH probes were designed using Biosearch Technologie’s Stellaris Probe Designer. Fluorophores used in this study were Quasar570 and Quasar670. For sample preparation, embryos were extruded from adult animals and subjected to freeze-crack on 0.01% poly-lysine coated slides followed by fixation in −20°C methanol for ≥ 15 minutes. Slides were washed five times in PBS with 0.1% Tween 20 (PBST) and fixed in 4% PFA in PBS for 1 hour at room temperature. Slides were again washed four times in PBST, twice in 2x SSC, and once in wash buffer (10% formamide, 2x SSC). Slides were then blocked in hybridization buffer (10% formamide, 2x SSC, 200 μg/mL BSA, 2mM Ribonucleoside Vanadyl Complex, 0.2 mg/mL yeast total RNA, 10% dextran sulfate) for 30 minutes at 37°C in a humid chamber. For hybridization, slides were incubated in 50-100 ⋂M probe in hybridization buffer at 37°C overnight. Slides were then washed twice in wash buffer at 37°C for 30 minutes, twice in 2x SSC, once in PBST and twice in PBS. Samples were mounted in ProLong Glass Antifade mountant and cured overnight.

### Combined *in situ* hybridization/immunofluorescence

Combined in situ hybridization with immunofluorescence was done by first doing the *in situ* protocol as described above. After the last wash in PBS, the slides were then re-fixed in 4% PFA for 1 hour at room temperature. The immunofluorescence protocol was then carried out as described above except 1 mg/mL UltraPure BSA (Thermo, cat# AM2616) was used in the blocking and antibody incubation steps. Primary antibody used: KT3 (DSHB) 1:100. Secondary antibody used: goat anti-mouse IgA conjugated to FITC (abcam, cat# ab97234) 1:500.

### Laser scanning confocal microscopy

Super-resolution microscopy was performed using a Zeiss LSM 880 microscope with a 63x-1.4 numerical aperture objective (Fig. 1, Fig. 3A, Fig. 4A, Fig. S1A,C, and Fig. S3A,B). The raw data was processed using default Airyscan settings with ZEN software. For Fig. 4A, representative high-resolution images were shown while the images used for quantification in Fig. 4B were collected by spinning disk confocal microscopy. All images shown are single Z slices.

### Spinning disk confocal microscopy

All other microscopy was performed using a Zeiss Axio Observer equipped with a CSU-W1 SoRA spinning disk scan head (Yokogawa). Images were taken using Slidebook software software (Intelligent Imaging Innovations) with a 63x objective with a 2.8x relay lens (Yokogawa). All images shown are single Z slices, except in Fig. 7C and D.

### Image quantification

All images were quantified in Fiji. For profile plots to show colocalization of granule components, a line was drawn through the center of a granule and the intensity along that line was measured using the plot profile tool in Fiji. Since the size of each granule varied slightly, the length of each plot was normalized to the smallest granule size. The intensities were then binned using the averageifs function in Excel. The background signal was subtracted and the intensities were normalized to the highest intensity.

For quantification of conditions that included sparse or asymmetrically localized RNAs/Proteins in P_4_, the sum intensity in P_4_ above threshold was measured and normalized to wild-type controls. The threshold was defined as being 1.5x the mean intensity of the entire embryo. To minimize background, the smooth function in Fiji was used, which replaces each pixel with the average of its 3×3 neighbors.

For quantification of symmetrically localized RNA/proteins in P_4_, the ratio of the mean intensity in the P blastomere over the mean intensity of a same sized region in the soma was measured. A background measurement was taken from outside the embryo and subtracted from the germline and soma intensities. The ratios were then normalized to wild type.

To assess the segregation of PGL-3 (Fig. S1D), DDX6^CGH-1^ and EDC-3 (Fig. S4) into P blastomeres, the mean intensity was measured in each P blastomere and were then normalized to the average P_0_ intensity.

To measure the ratio of RNA inside/outside of granules, the granule (labeled by MEG-1::GFP in Fig. 3B, SL1 in Fig. S5 or PGL-3 in Fig. S7) was defined as being 1.5x above the mean intensity of the signal within the P blastomere. The mean intensity inside and outside the granule in the cytoplasm was measured. A background signal was taken from a region outside the embryo and subtracted.

### Statistical analysis and plotting

Perseus (Tyanova et al., 2016b) was used for t-tests on mass spec data. To determine the significance of the enrichment of P-body proteins in MEG-1 immunoprecipitates, we assumed a total pool of 6,000 proteins, which is roughly the size of the embryonic proteome (Saha et al., 2016).

Statistics for differential expression analysis were done using DESeq2 (Love et al., 2014). To determine the significance of the overlap between predicted POS-1 targets (Elewa et al., 2015) and *meg-1 meg-2* differentially expressed genes, we assumed a total pool of 11,121 transcripts. We arrived at this number by setting an FPKM threshold in our RNA-seq analysis of 0.002178852 FPKM, which was the lowest FPKM in *meg-1 meg-2* animals that we were able to detect a significant increase in gene expression. Any non-protein coding genes were also identified and removed from the list by using the SimpleMine tool on WormBase.

All other statistical analysis was conducted using R or Graphpad Prism 9 software. Data were plotted with either Graphpad Prism 9 or ggplot2 (Wickham, 2016).

---

**Supplemental table 1:** Sterility of *meg-1 meg-2(ax4532)* mutant.

| Genotype | percent sterile (n) |
| --- | --- |
| MEG-1::GFP/*meg-1 meg-2 (ax4532)* (+/-) | 0 (289) |
| *meg-1 meg-2 (ax4532)/meg-1 meg-2 (ax4532)* (-/-) | 100 (156) |

**Supplemental table 4:**
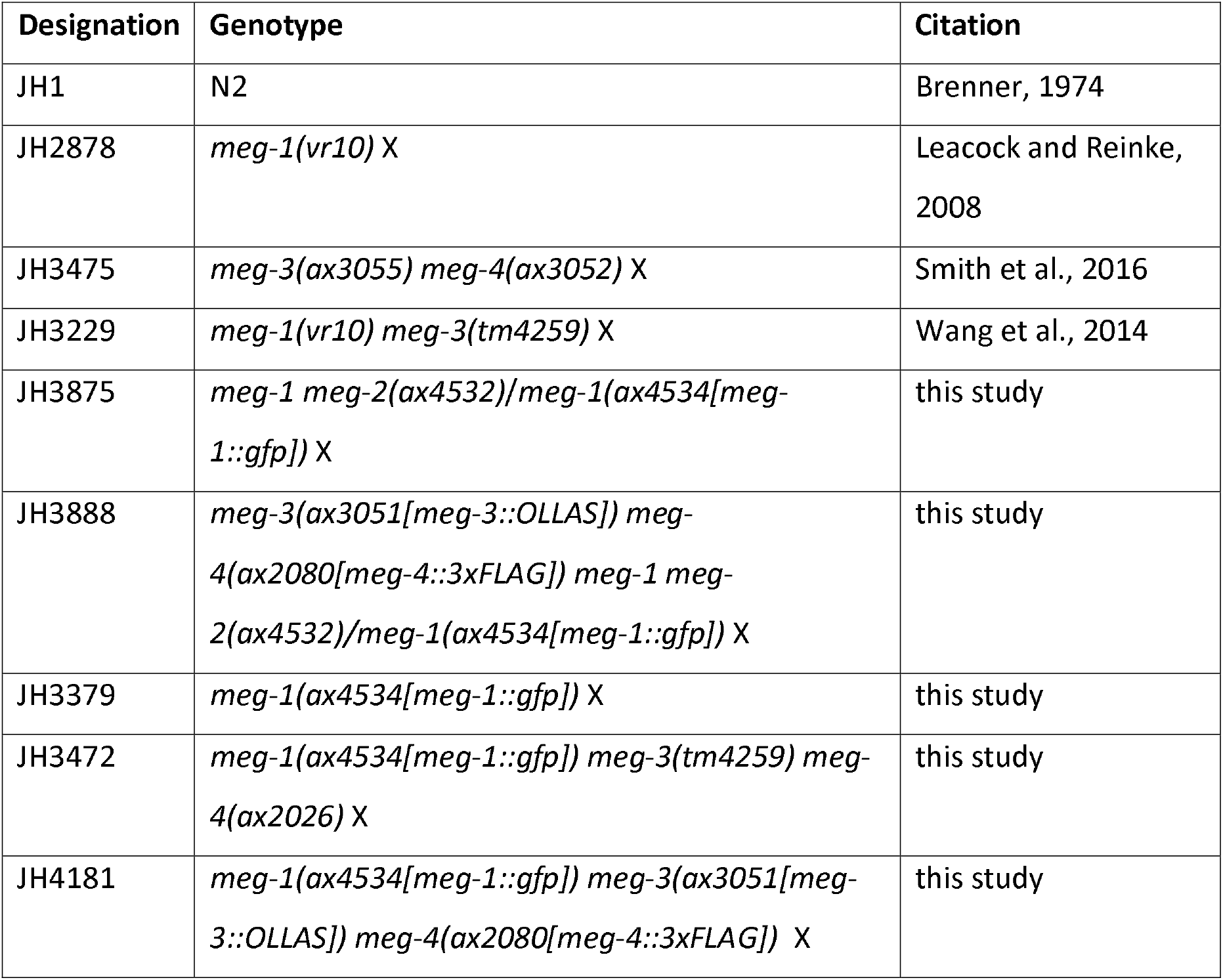
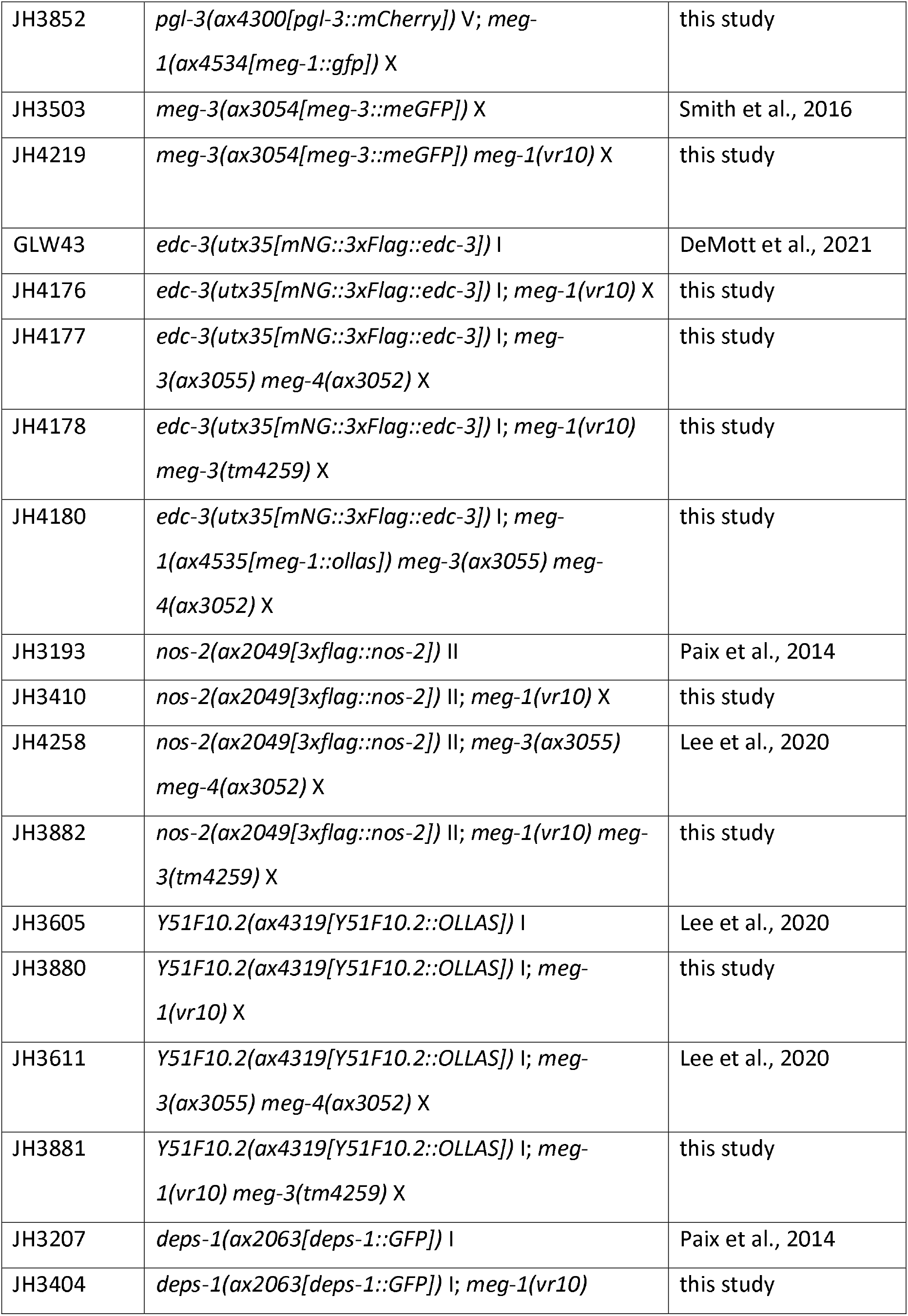
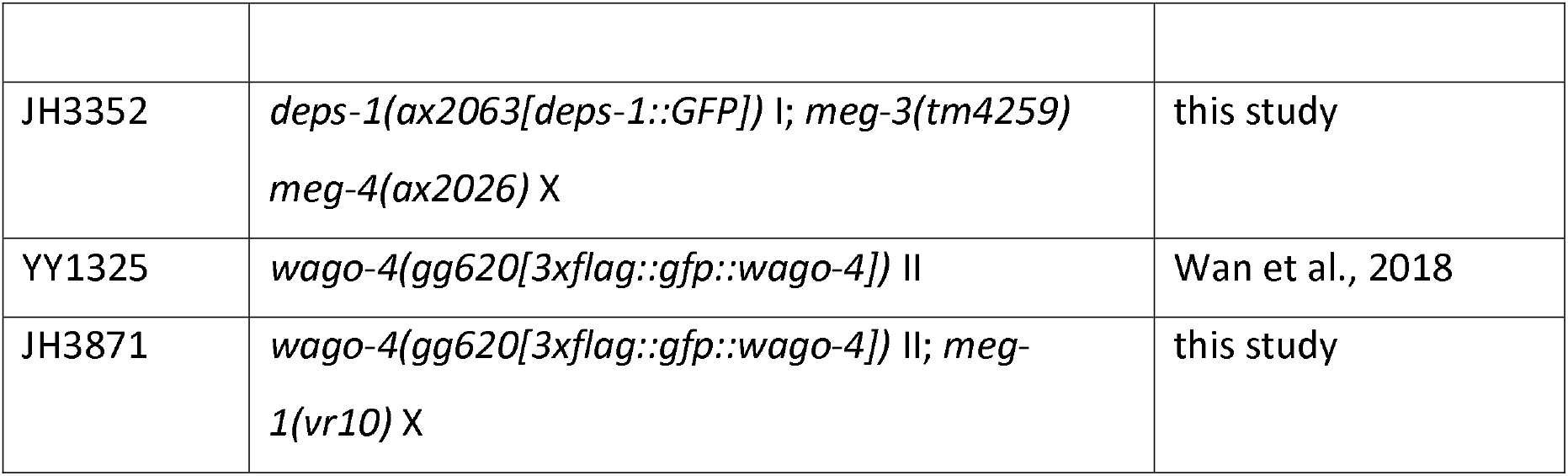
Strains used.

| Designation | Genotype | Citation |
| --- | --- | --- |
| JH1 | N2 | Brenner, 1974 |
| JH2878 | *meg-1(vr10)* X | Leacock and Reinke, 2008 |
| JH3475 | *meg-3(ax3055) meg-4(ax3052)* X | Smith et al., 2016 |
| JH3229 | *meg-1(vr10) meg-3(tm4259)* X | Wang et al., 2014 |
| JH3875 | *meg-1 meg-2(ax4532)/meg-1(ax4534[meg-1::gfp])* X | this study |
| JH3888 | *meg-3(ax3051[meg-3::OLLAS]) meg-4(ax2080[meg-4::3xFLAG]) meg-1 meg-2(ax4532)/meg-1(ax4534[meg-1::gfp])* X | this study |
| JH3379 | *meg-1(ax4534[meg-1::gfp])* X | this study |
| JH3472 | *meg-1(ax4534[meg-1::gfp]) meg-3(tm4259) meg-4(ax2026)* X | this study |
| JH4181 | *meg-1(ax4534[meg-1::gfp]) meg-3(ax3051[meg-3::OLLAS]) meg-4(ax2080[meg-4::3xFLAG])* X | this study |
| JH3852 | *pgl-3(ax4300[pgl-3::mCherry])V; meg-1(ax4534[meg-1::gfp]) X* | this study |
| JH3503 | *meg-3(ax3054[meg-3::meGFP]) X* | Smith et al., 2016 |
| JH4219 | *meg-3(ax3054[meg-3::meGFP]) meg-1(vr10) X* | this study |
| GLW43 | *edc-3(utx35[mNG::3xFlag::edc-3]) I* | DeMott et al., 2021 |
| JH4176 | *edc-3(utx35[mNG::3xFlag::edc-3]) I; meg-1(vr10) X* | this study |
| JH4177 | *edc-3(utx35[mNG::3xFlag::edc-3]) I; meg-3(ax3055) meg-4(ax3052) X* | this study |
| JH4178 | *edc-3(utx35[mNG::3xFlag::edc-3]) I; meg-1(vr10) meg-3(tm4259) X* | this study |
| JH4180 | *edc-3(utx35[mNG::3xFlag::edc-3]) I; meg-1(ax4535[meg-1::ollas]) meg-3(ax3055) meg-4(ax3052) X* | this study |
| JH3193 | *nos-2(ax2049[3xflag::nos-2]) II* | Paix et al., 2014 |
| JH3410 | *nos-2(ax2049[3xflag::nos-2]) II; meg-1(vr10) X* | this study |
| JH4258 | *nos-2(ax2049[3xflag::nos-2]) II; meg-3(ax3055) meg-4(ax3052) X* | Lee et al., 2020 |
| JH3882 | *nos-2(ax2049[3xflag::nos-2]) II; meg-1(vr10) meg-3(tm4259) X* | this study |
| JH3605 | *Y51F10.2(ax4319[Y51F10.2::OLLAS]) I* | Lee et al., 2020 |
| JH3880 | *Y51F10.2(ax4319[Y51F10.2::OLLAS]) I; meg-1(vr10) X* | this study |
| JH3611 | *Y51F10.2(ax4319[Y51F10.2::OLLAS]) I; meg-3(ax3055) meg-4(ax3052) X* | Lee et al., 2020 |
| JH3881 | *Y51F10.2(ax4319[Y51F10.2::OLLAS]) I; meg-1(vr10) meg-3(tm4259) X* | this study |
| JH3207 | *deps-1(ax2063[deps-1::GFP]) I* | Paix et al., 2014 |
| JH3404 | *deps-1(ax2063[deps-1::GFP]) I; meg-1(vr10)* | this study |
| JH3352 | *deps-1(ax2063[deps-1::GFP])* l; *meg-3(tm4259)*<br>*meg-4(ax2026)* X | this study |
| YY1325 | *wago-4(gg620[3xflag::gfp::wago-4])* ll | Wan et al., 2018 |
| JH3871 | *wago-4(gg620[3xflag::gfp::wago-4])* ll; *meg-1(vr10)* X | this study |

