## supplemental figures for "Specialized germline P-bodies are required to specify germ cell fate in *C. elegans* embryos"

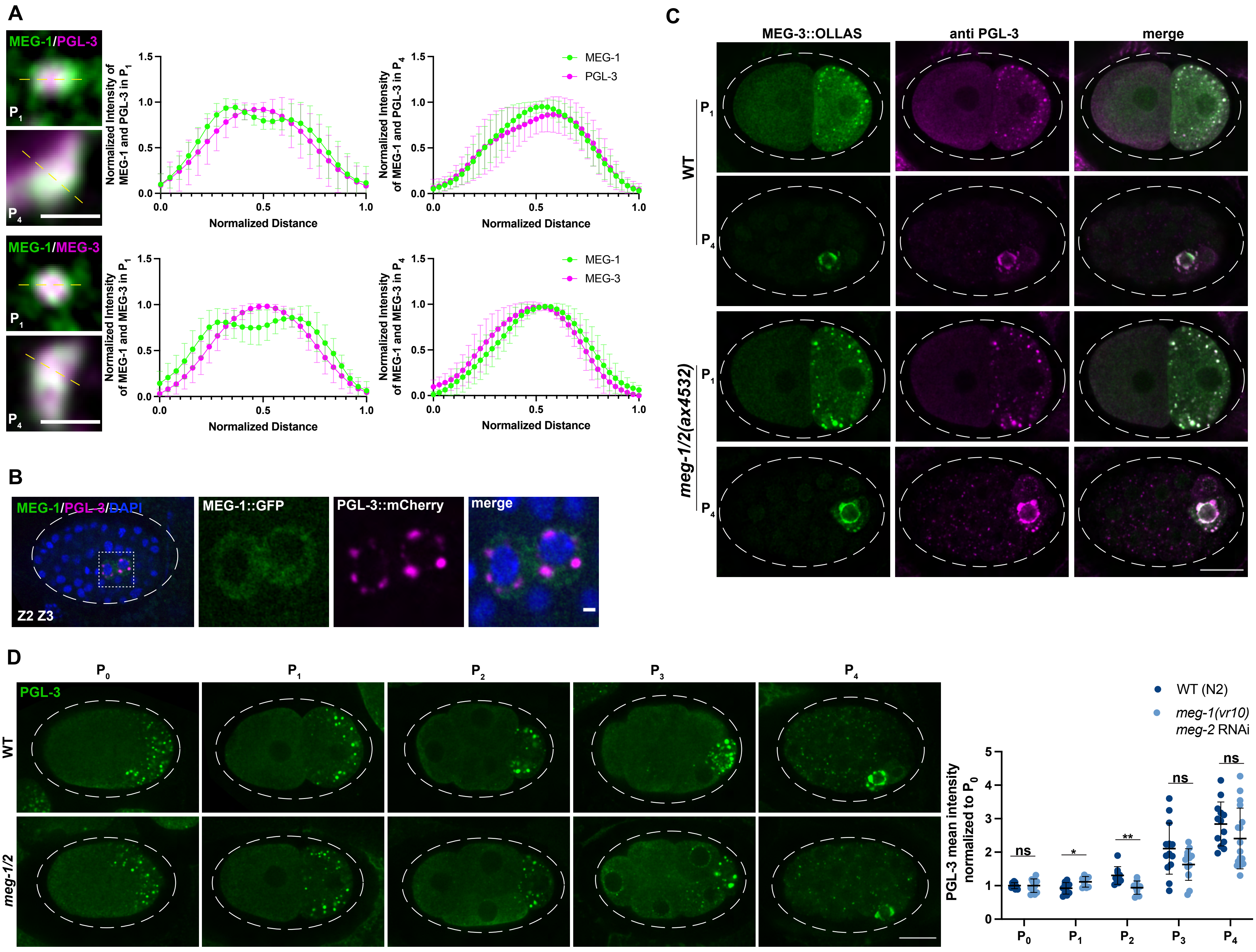
A
MEG-1/PGL-3
MEG-1/MEG-3
Normalized Intensity of MEG-1 and PGL-3 in P1
Normalized Intensity of MEG-1 and PGL-3 in P4
Normalized Intensity of MEG-1 and MEG-3 in P1
Normalized Intensity of MEG-1 and MEG-3 in P4
Normalized Distance
MEG-1
PGL-3
MEG-3
B
MEG-1/PGL-3/DAPI
MEG-1::GFP
PGL-3::mCherry
merge
Z2 Z3
C
MEG-3::OLLAS
anti PGL-3
merge
WT
meg-1/2(ax4532)
D
PGL-3
WT
meg-1/2
WT (N2)
meg-1(vr10) meg-2 RNAi
PGL-3 mean intensity normalized to P0
ns
*
**

**Fig S1: MEG-1 puncta are distinct from P granules.**

(A) Airyscan photomicrographs of granules from embryos expressing MEG-1::GFP and co-stained for GFP and PGL-3 or MEG-3::OLLAS. Dashed yellow lines indicate where intensity measurements were made for quantification. Plots show relative intensity through the center of the granules of MEG-1 and PGL-3 in $P_1$ (n=12 granules from 2 embryos) and $P_4$ (n=9 granules from 2 embryos) and MEG-1 and MEG-3 in $P_1$ (n=17 granules from 2 embryos) and $P_4$ (n=8 granules from 2 embryos). Error bars represent mean ± s.d. Scale bars are 1 µm. (B) Representative photomicrograph of a ~100-cell embryo expressing MEG-1::GFP and PGL-3::mCherry. Z2 and Z3 are shown in inset. At this stage, MEG-1 becomes dispersed in the cytoplasm, whereas PGL-3 remains in perinuclear granules. Scale bar is 1 µm. (C) Representative Airyscan photomicrographs of the indicated genotypes expressing MEG-3::OLLAS and co-stained for OLLAS and PGL-3. P granule assembly and segregation to germline blastomeres is not dependent on *meg-1 meg-2*. Scale bar is 10 µM. (D) Representative photomicrographs of embryos of the indicated genotype stained for PGL-3. In this and all subsequent figures, the designation *meg-1/2* refers to *meg-1(vr10) meg-2* RNAi. PGL-3 mean intensity in P blastomeres was normalized to $P_0$. On average, PGL-3 accumulation into $P_2$ was less efficient in *meg-1 meg-2* embryos, but was not significantly different by the $P_4$ stage. Scale bar is 10 µm. Data shown are from two independent experiments where mutant and control animals were processed in parallel. Number of embryos quantified per stage for WT: $P_0$=9, $P_1$= 10, $P_2$= 9, $P_3$=14, $P_4$=13. For *meg-1/2*: $P_0$=11, $P_1$=8, $P_2$=9, $P_3$=13, $P_4$=17. ***P*≤0.01, **P*≤0.05, ns=not significant (*t*-test). Error bars represent mean ± s.d.

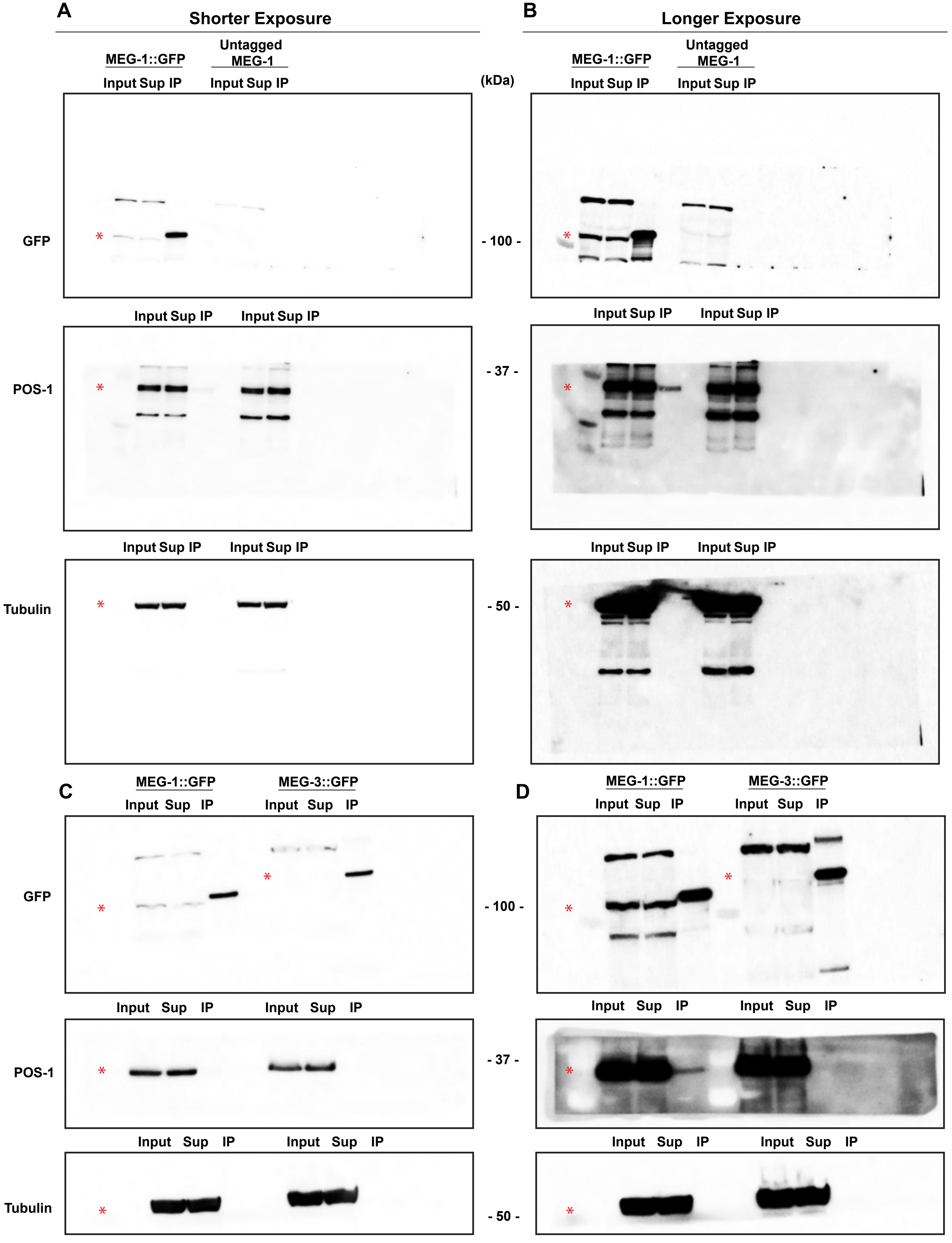
A
Shorter Exposure
MEG-1::GFP
Untagged MEG-1
Input Sup IP
Input Sup IP
(kDa)
GFP
- 100 -
POS-1
- 37 -
Tubulin
- 50 -
B
Longer Exposure
MEG-1::GFP
Untagged MEG-1
C
MEG-1::GFP
MEG-3::GFP
Input Sup IP
D
*

**Fig S2: Western blots of MEG-1::GFP immunoprecipitations.**

Full western blot images from immunoprecipitation experiment shown in Fig. 3B (A and B) and 3C (C and D). Shorter exposures are shown in (A) and (C) and longer exposures of the same blot are shown in (B) and (D). Red asterisks indicate where the expected protein band would migrate. All other bands on the blots are non-specific.

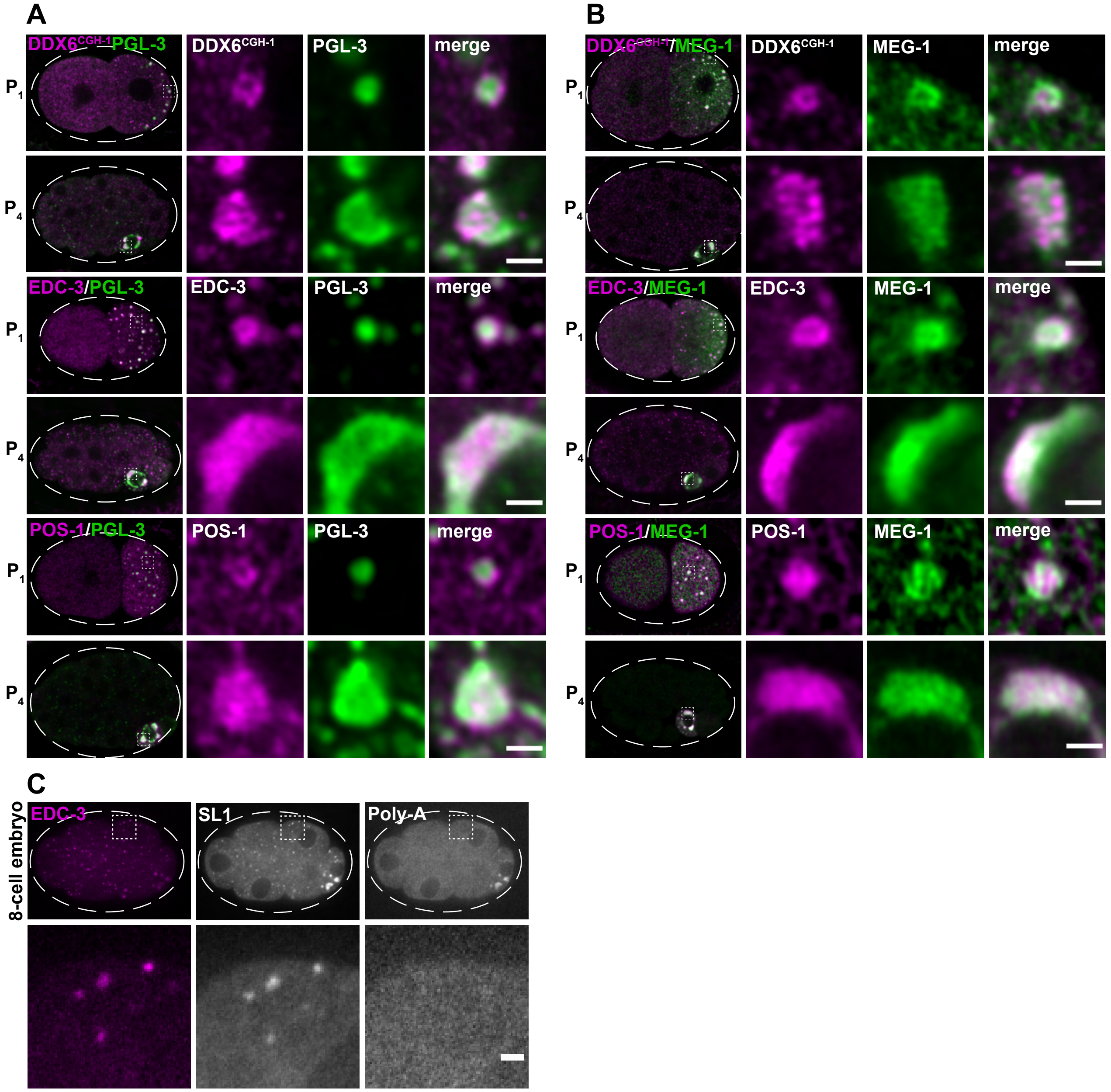


**Fig S3: Distribution of germline P-body proteins.**

(A) Airyscan photomicrographs of embryos either stained for DDX6$^{CGH\text{-}1}$, expressing mNG::3xFLAG::EDC-3, or stained for POS-1 and co-stained for PGL-3. Insets show granules in $P_1$ and $P_4$. In $P_1$, P-body proteins are enriched at the periphery of PGL-3. In $P_4$, they overlap with PGL-3. (B) Airyscan photomicrographs of embryos expressing MEG-1::GFP and either stained for DDX6$^{CGH\text{-}1}$, expressing mNG::3xFLAG::EDC-3, or stained for POS-1. Insets show granules in $P_1$ and $P_4$. In $P_1$, MEG-1 and P-body proteins form complex partially overlapping patterns. In $P_4$, they overlap. (C) Representative photomicrograph of 8-cell embryo expressing mNeonGreen::3xFLAG::EDC-3 and probed for SL1 and poly-A. Note that P granules (right most cell) are positive for both. Inset shows a somatic cell where EDC-3 foci are enriched for SL1, but not poly-A. Scale bars are 1 µm.

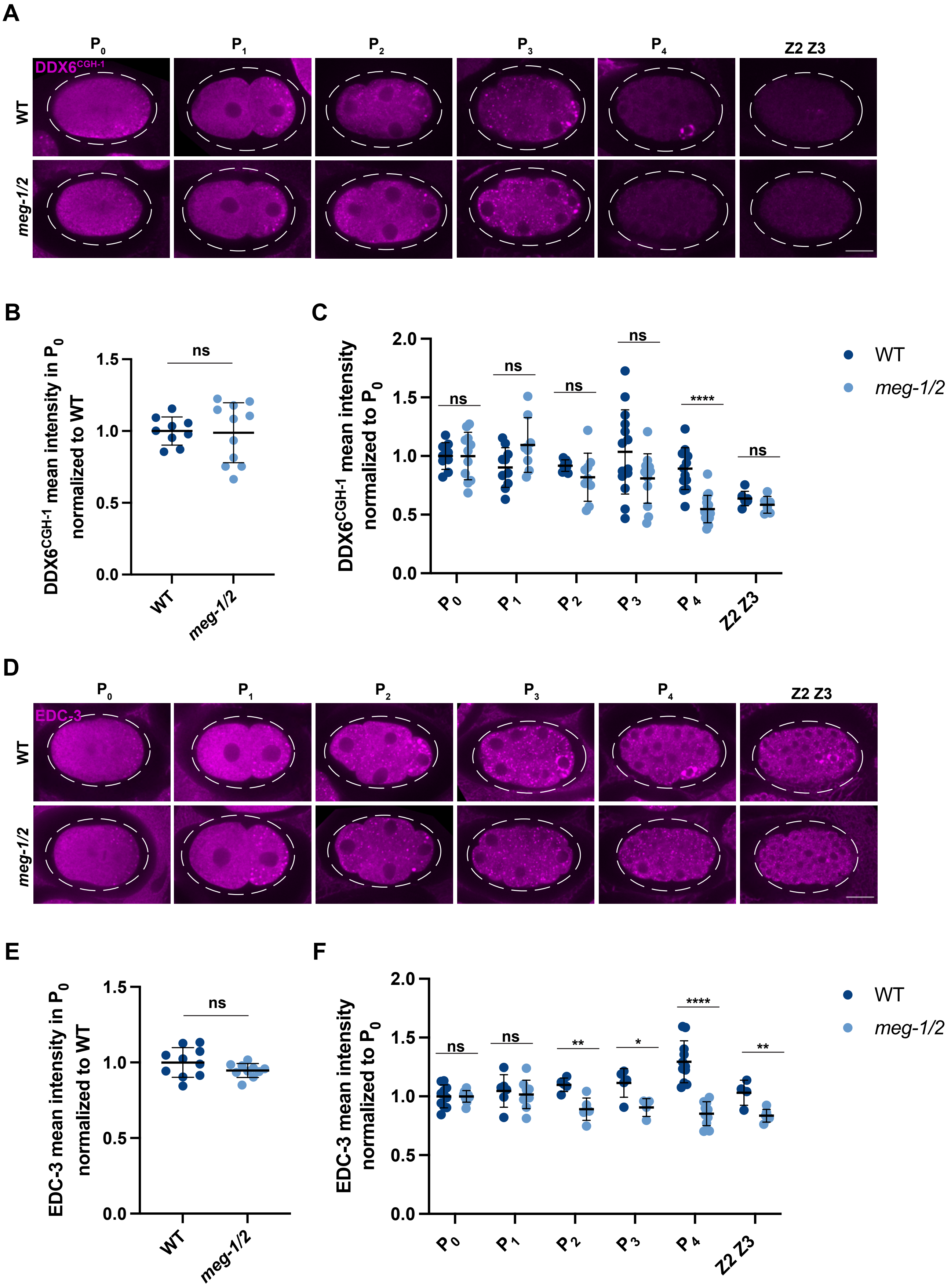
A
P0
P1
P2
P3
P4
Z2 Z3
DDX6CGH-1
WT
meg-1/2
B
DDX6CGH-1 mean intensity in P0 normalized to WT
ns
1.5
1.0
0.5
0.0
WT
meg-1/2
C
DDX6CGH-1 mean intensity normalized to P0
2.0
1.5
1.0
0.5
0.0
ns
ns
ns
ns
****
ns
P0
P1
P2
P3
P4
Z2 Z3
WT
meg-1/2
D
P0
P1
P2
P3
P4
Z2 Z3
EDC-3
WT
meg-1/2
E
EDC-3 mean intensity in P0 normalized to WT
ns
1.5
1.0
0.5
0.0
WT
meg-1/2
F
EDC-3 mean intensity normalized to P0
2.0
1.5
1.0
0.5
0.0
ns
ns
**
*
****
**
P0
P1
P2
P3
P4
Z2 Z3
WT
meg-1/2

**Fig S4. *meg-1* and *meg-2* are required to stabilize $DDX6^{CGH-1}$ and EDC-3 in $P_4$.**
(A) Photomicrographs of embryos of the indicated genotypes stained for $DDX6^{CGH-1}$. $DDX6^{CGH-1}$ distribution is unchanged in *meg-1 meg-2* embryos compared to wild-type up to the 8-cell stage. After the 8-cell stage, $DDX6^{CGH-1}$ is turned over in somatic blastomeres in wild-type, and in all cells in *meg-1 meg-2* embryos. Shortly after the 100-cell stage, $DDX6^{CGH-1}$ is uniformly present in low concentrations in both genotypes. (B) Total levels of maternally provided $DDX6^{CGH-1}$ to $P_0$ zygotes are the same in *meg-1 meg-2* embryos compared to wild-type. (C) Mean intensity of $DDX6^{CGH-1}$ in each P blastomere normalized to $P_0$. Quantification for each genotype is from two independent experiments where mutant and control animals were processed in parallel. Number of embryos quantified per stage for WT: $P_0$=9, $P_1$=10, $P_2$=9, $P_3$=14, $P_4$=13, Z2 Z3=7. For *meg-1/2*: $P_0$=11, $P_1$=8, $P_2$=9, $P_3$=13, $P_4$=17, Z2 Z3=6. (D) Photomicrographs of embryos of the indicated genotypes expressing mNG::3xFLAG::EDC-3 and stained for FLAG. Beginning in $P_2$, EDC-3 distribution is lower in *meg-1 meg-2* embryos compared to wild-type and is most dramatically reduced in $P_4$. (E) Total levels of maternally provided EDC-3 to $P_0$ zygotes are the same in *meg-1 meg-2* embryos compared to wild-type. (F) Mean intensity of EDC-3 in each P blastomere normalized to $P_0$. Quantification for each genotype is from one experiment where mutant and control animals were processed in parallel. Number of embryos quantified per stage for WT: $P_0$=10, $P_1$=6, $P_2$=5, $P_3$=5, $P_4$=11, Z2 Z3=4. For *meg-1/2*: $P_0$=10, $P_1$=9, $P_2$=6, $P_3$=4, $P_4$=10, Z2 Z3=6. ****$P \leq 0.0001$, **$P \leq 0.01$, *$P \leq 0.05$, ns=not significant (*t*-test). All error bars represent mean ± s.d. All scale bars are 10 μm.

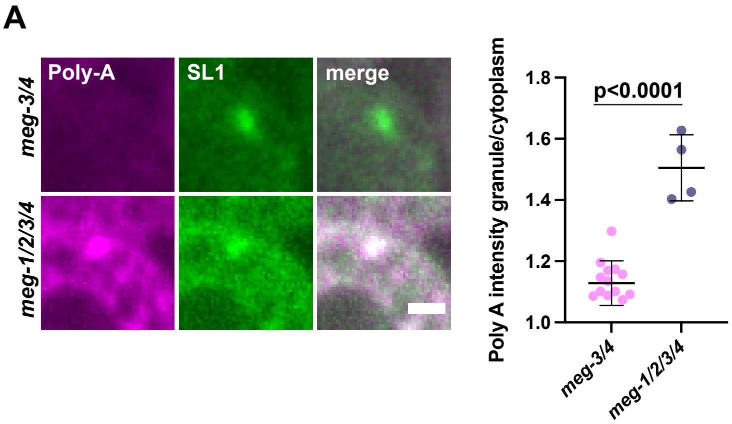


**Fig S5: Distribution of poly-A in SL1 puncta.**

(A) Photomicrographs of SL1 foci in $P_4$ of the indicated genotypes. SL1 foci do not enrich poly-A in *meg-3 meg-4* embryos, but do in *meg-1 meg-2 meg-3 meg-4*. Embryos quantified are from the experiment shown in Fig. 4D, but only including embryos that had SL1 foci in the center z-plane of $P_4$. For *meg-3/4* n=14, for *meg-1/2/3/4* n=4. Scale bar is 1 µm. A *t*-test was used to make comparisons between genotypes. Error bars represent mean ± s.d.

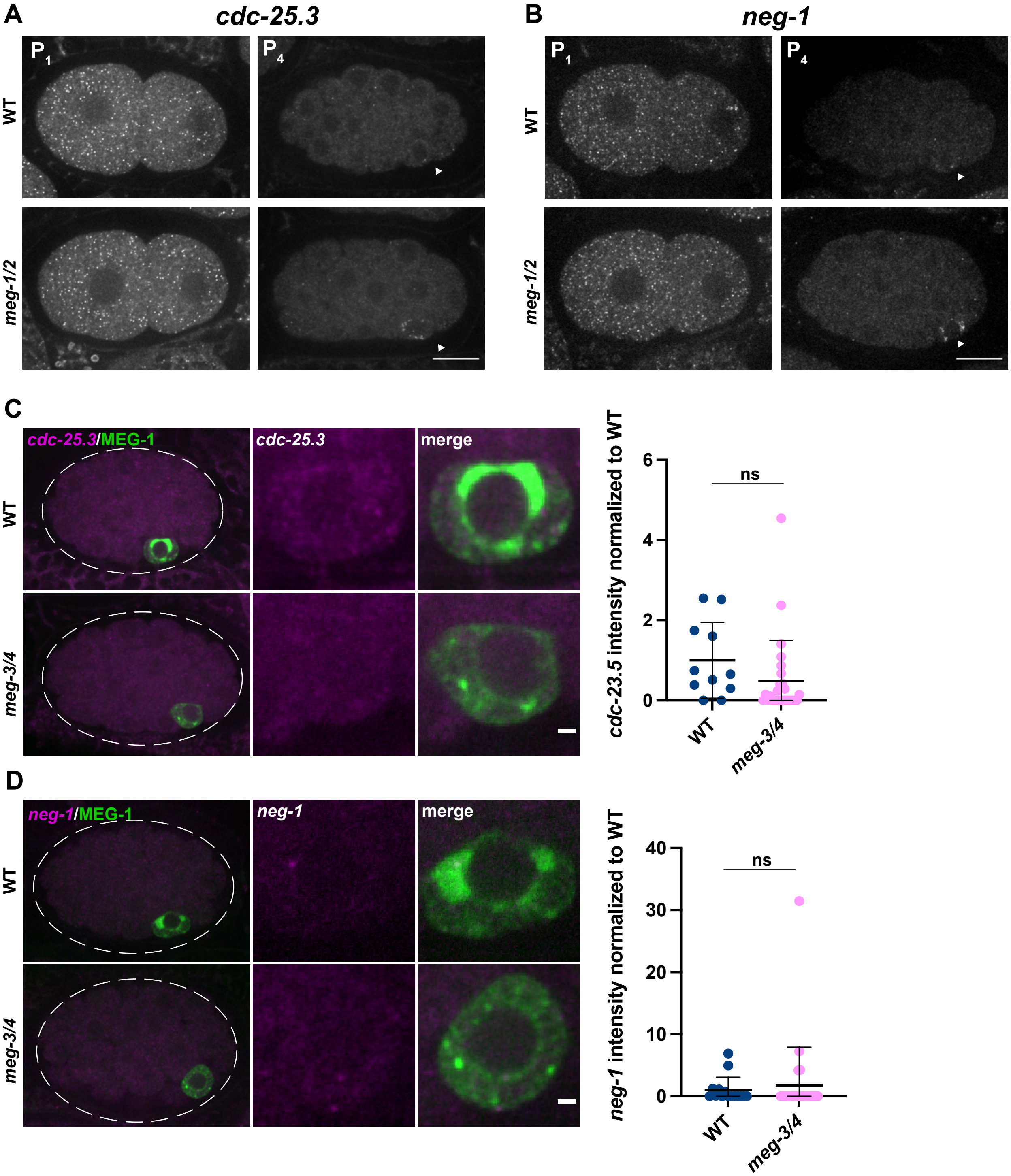
A
cdc-25.3
B
neg-1
P1
P4
WT
meg-1/2
C
cdc-25.3/MEG-1
cdc-25.3
merge
meg-3/4
cdc-23.5 intensity normalized to WT
ns
6
4
2
0
D
neg-1/MEG-1
neg-1
neg-1 intensity normalized to WT
40
30
20
10

**Fig S6: Spatiotemporal localization of *cdc-25.3* and *neg-1.***

Representative photomicrographs of embryos of the indicated genotypes probed for *cdc-25.3* (A) and *neg-1* (B). Both transcripts are maternally deposited and mostly turned over by the 28-cell stage, except in $P_4$ in *meg-1 meg-2* embryos. Arrows point to $P_4$. Scale bars are 10 µm. (C) Representative photomicrographs of embryos of the indicated genotypes expressing MEG-1::GFP and probed for *cdc-25.3.* Embryos were quantified from two independent experiments where mutant and control animals were processed in parallel. Wild type n=11, *meg-3/4* n=26. (D) Representative photomicrographs of embryos of the indicated genotypes expressing MEG-1::GFP and probed for *neg-1*. Embryos were quantified from two independent experiments where mutant and control animals were processed in parallel. Wild type n=15, *meg-3/4* n=27. *cdc-25.3* and *neg-1* are efficiently turned over in *meg-3 meg-4* embryos. Scale bars in (C) and (D) are 1 µm. Error bars represent mean ± s.d. A *t*-test was used to make comparisons between genotypes.

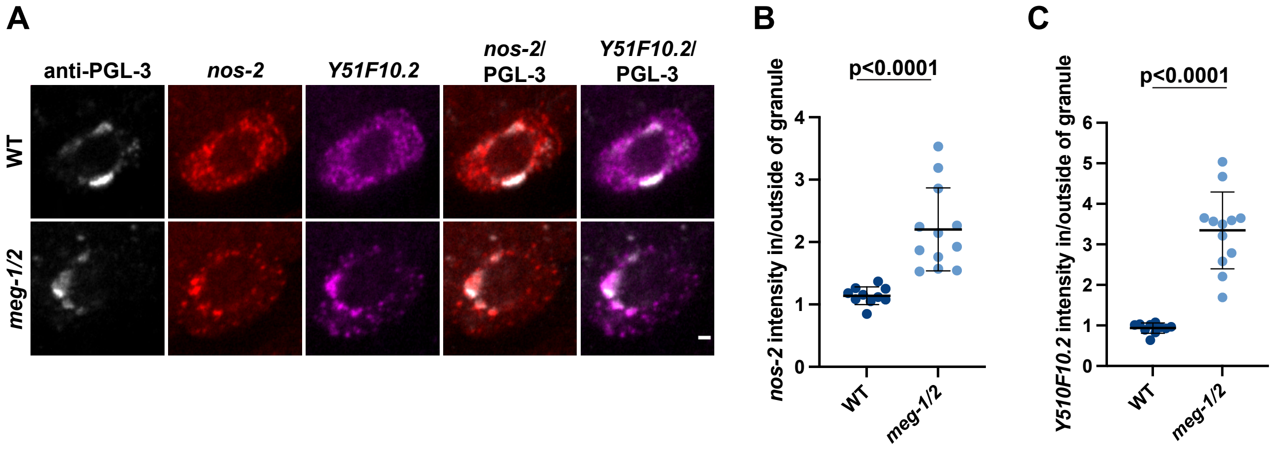


**Fig S7: *nos-2* and *Y51F10.2* RNAs are enriched in P granules in *meg-1 meg-2* $P_4$ blastomeres.**

(A) Photomicrographs of $P_4$ in the indicated genotypes stained for PGL-3 and probed for *nos-2* and *Y51F10.2*. Scale bar is 1 μm. The ratio of *nos-2* (B) and *Y51F10.2* (C) inside vs. outside of the PGL-3 granule is quantified. In *meg-1 meg-2* embryos, these transcripts are more enriched in the granule compared to wild-type. Quantification for each genotype is from one experiment where mutant and control animals were processed in parallel. Number of embryos quantified for wild type=10; for *meg-1/2*=12. A *t*-test was used to make comparisons between genotypes. Error bars represent mean ± s.d.

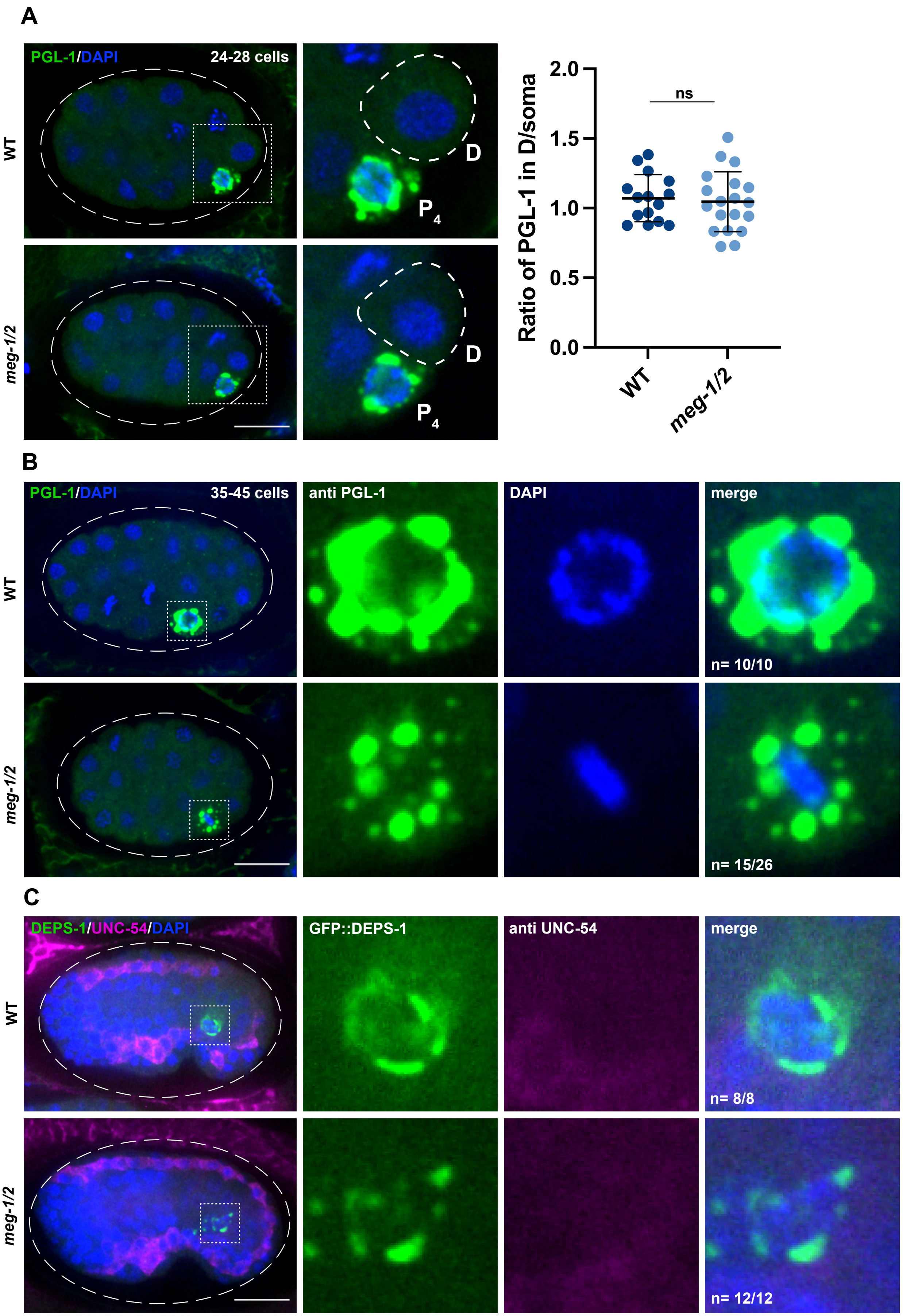
A
PGL-1/DAPI
24-28 cells
WT
meg-1/2
D
P4
Ratio of PGL-1 in D/soma
2.0
1.5
1.0
0.5
0.0
ns
WT
meg-1/2
B
PGL-1/DAPI
35-45 cells
anti PGL-1
DAPI
merge
n= 10/10
n= 15/26
C
DEPS-1/UNC-54/DAPI
GFP::DEPS-1
anti UNC-54
merge
n= 8/8
n= 12/12

**Fig S8: Behavior of germ cells in *meg-1 meg-2* embryos.**

(A) Photomicrographs of 24-28 cell stage embryos of the indicated genotypes stained for PGL-1. Inset shows $P_4$ and its sister cell, D. The ratio of PGL-1 intensity in D over a neighboring somatic cell was not different in *meg-1 meg-2* embryos compared to wild-type, indicating that PGL-1 is properly asymmetrically distributed to $P_4$ in *meg-1 meg-2* embryos. Quantification for each genotype is from one experiment where mutant and control animals were processed in parallel. Number of embryos quantified for wild type=15; for *meg-1/2*=19. A *t*-test was used to make comparisons between genotypes. Error bars represent mean ± s.d. (B) Photomicrographs of 35-45 cell stage embryos of the indicated genotypes stained for PGL-1. Inset shows $P_4$. 10/10 wild-type embryos had characteristic chromatin organization in $P_4$, whereas in 15/26 *meg-1 meg-2* embryos, $P_4$ began to divide precociously, indicated by chromatin in metaphase (as shown) or other stages of mitosis. (C) Photomicrographs of bean stage embryos of the indicated genotypes expressing DEPS-1::GFP and stained for myosin heavy chain (UNC-54). PGL-1 positive cells in *meg-1 meg-2* embryos do not express myosin despite transcribing *hlh-1* (Fig 7). All scale bars are 10 μm.
